# High-throughput discovery and deep characterization of cyclin-CDK docking motifs

**DOI:** 10.1101/2024.12.03.625240

**Authors:** Mihkel Örd, Matthew J. Winters, Mythili S. Subbanna, Natalia de Martin Garrido, Victoria I. Cushing, Johanna Kliche, Caroline Benz, Ylva Ivarsson, Basil J. Greber, Peter M. Pryciak, Norman E. Davey

**Affiliations:** University of Cambridge, CRUK Cambridge Institute, Cambridge, UK; The Institute of Cancer Research, Chester Beatty Laboratories, 237 Fulham Road, London, SW3 6JB, UK; Department of Biochemistry and Molecular Biotechnology, University of Massachusetts Chan Medical School, Worcester, MA 01605, USA; Department of Chemistry - BMC, Uppsala University, Husargatan 3, 751 23 Uppsala, Sweden

**Keywords:** kinase, cyclin, cyclin-dependent kinases, short linear motif, SLiMs, enzyme docking, binding specificity, binding affinity, phosphorylation, post-translational modification

## Abstract

Cyclin-CDKs are master regulators of cell division. In addition to directly activating the CDK, the cyclin subunit regulates CDK specificity by binding short peptide “docking” motifs in CDK substrates. Here, we measure the relative binding strength of ∼100,000 peptides to 11 human cyclins from five cyclin families (D, E, A, B and F). Using a quantitative intracellular binding assay and large-scale tiled peptide screening, we identified a range of non-canonical binders that unveil a broader than anticipated repertoire of cyclin docking motif types. Structural and saturation mutagenesis studies revealed distinct binding modes and sequence features that govern motif recognition, binding strength, and cyclin preference. Docking motifs vary from highly selective to pan-cyclin, thereby fine-tuning the timing of CDK phosphorylation during cell cycle progression. Overall, these findings provide an unprecedented depth of understanding about the rules encoding specificity and affinity within a group of related but distinct protein domains.

## Introduction

Cell division requires precise orchestration of diverse molecular processes, such as DNA replication, chromosome segregation, the centrosome cycle, transcription and metabolism^1^. Each step is coordinated by cyclin-dependent kinases (CDKs) that phosphorylate hundreds of proteins in a temporally resolved manner during the cell cycle^2–5^. The key determinant of these cycles of phosphorylation is the tightly controlled expression, localization, and degradation of cyclins^1^. In metazoans, four major classes of cyclin-CDK complexes are sequentially activated during the cell cycle, with cyclin D-Cdk4/6 and E-Cdk2 initiating cell cycle entry, cyclin A-Cdk1/2 driving S phase and mitotic entry, and cyclin B-Cdk1 coordinating mitosis^6^. In G1 phase, CDK activity is minimal due to the expression of CDK inhibitory proteins (e.g., p27) and the low levels of cyclins E, A and B. The accumulation of CDK activity that peaks in mitosis is driven by the expression of cyclins together with inactivation of their inhibitory proteins, and the gradually increasing intrinsic activity of subsequent cyclin-CDK complexes^1,7,8^. Finally, the system is reset by the degradation of cyclins A and B during mitosis, and the subsequent dephosphorylation of CDK substrates^9^.

Several mechanisms control the phosphorylation of CDK targets. First, CDKs are serine/threonine kinases that phosphorylate minimal [ST]P and full [ST]Px[KR] consensus sites, though Cdk1 can also phosphorylate sites lacking the adjacent proline^10,11^. Second, substrate multisite phosphorylation can be promoted by the CDK-bound Cks1 subunit that binds pre-phosphorylated TP sites^10,12–14^. Third, cyclin subunits recruit specific proteins by binding to docking motifs^15^. In particular, “RxL” docking motifs are recognized by a binding pocket called the hydrophobic patch that is conserved in all major classes of cell cycle cyclins (D, E, A, B)^16,17^. In CDK substrates, RxL motifs enhance phosphorylation, whereas in CDK inhibitors they block substrate recognition^18,19^. Moreover, these interactions can control cyclin localization in human and yeast cells^20–22^. Cyclin F also recognizes RxL motifs but does not bind CDKs, acting instead as a substrate recognition subunit of the SCF (Skp1–Cul1–F-box protein) ubiquitin ligase complex^23^. Additionally, cyclin D docks to C-terminal helices in Rb-family proteins^8^, and cyclin B binds to a helix in Mad1 and a phosphoserine-containing motif in separase^24–26^. Further diversity of cyclin docking has been characterized in budding yeast, where specific docking motifs have been found for most cyclin types^7,27–30^.

Over 30 human proteins have been found to recruit cyclins through RxL docking motifs. These motifs generally conform to a strict consensus where an invariant leucine (P0, which we refer to as Φ_0_) is flanked N-terminally by an arginine or lysine residue (P-2) and C-terminally by a phenylalanine or a leucine (at P+1 or P+2, which we refer to as Φ_C_) (**Fig 1A-B**). Co-crystal structures of cyclin A2 with RxL peptides explain these sequence preferences^31–33^, revealing that the two consensus hydrophobic residues are buried in a hydrophobic pocket and the N-terminal portion of the peptide makes extensive contacts with the cyclin surface anchored around the consensus basic position (**Fig 1C**). Distinct backbone conformations permit the Φ_C_ hydrophobic residue to exist at either the P+1 or P+2 position. A handful of atypical motifs have been identified, suggesting that additional RxL-like motifs exist. For example, a peptide from SAMHD1 conforms to the hydrophobic positions but lacks a basic residue at P-2^34^, while a motif in RRM2 that binds Cyclin F features a P0 Ile^35^. Moreover, SKP2 contains two atypical motifs, one that binds the RxL-binding surface in a reverse orientation, and one that binds a distinct cyclin surface^36,37^. The cyclin residues contacting the RxL motif exhibit sequence diversity across the human cyclins, particularly in the residues contacting the N-terminal portion of the motif (**Fig 1D**). Despite this, relatively little is known about the tolerated residues in RxL motifs, how the motif sequence context can modulate binding strength, whether different cyclins show any distinct sequence preferences, or what are the targets of specific cyclins. Given the critical role of cyclin-CDKs in cell proliferation and the considerable interest in inhibiting these complexes for therapeutic purposes^38,39^, a systematic effort to address these questions is warranted.

**Figure 1.**
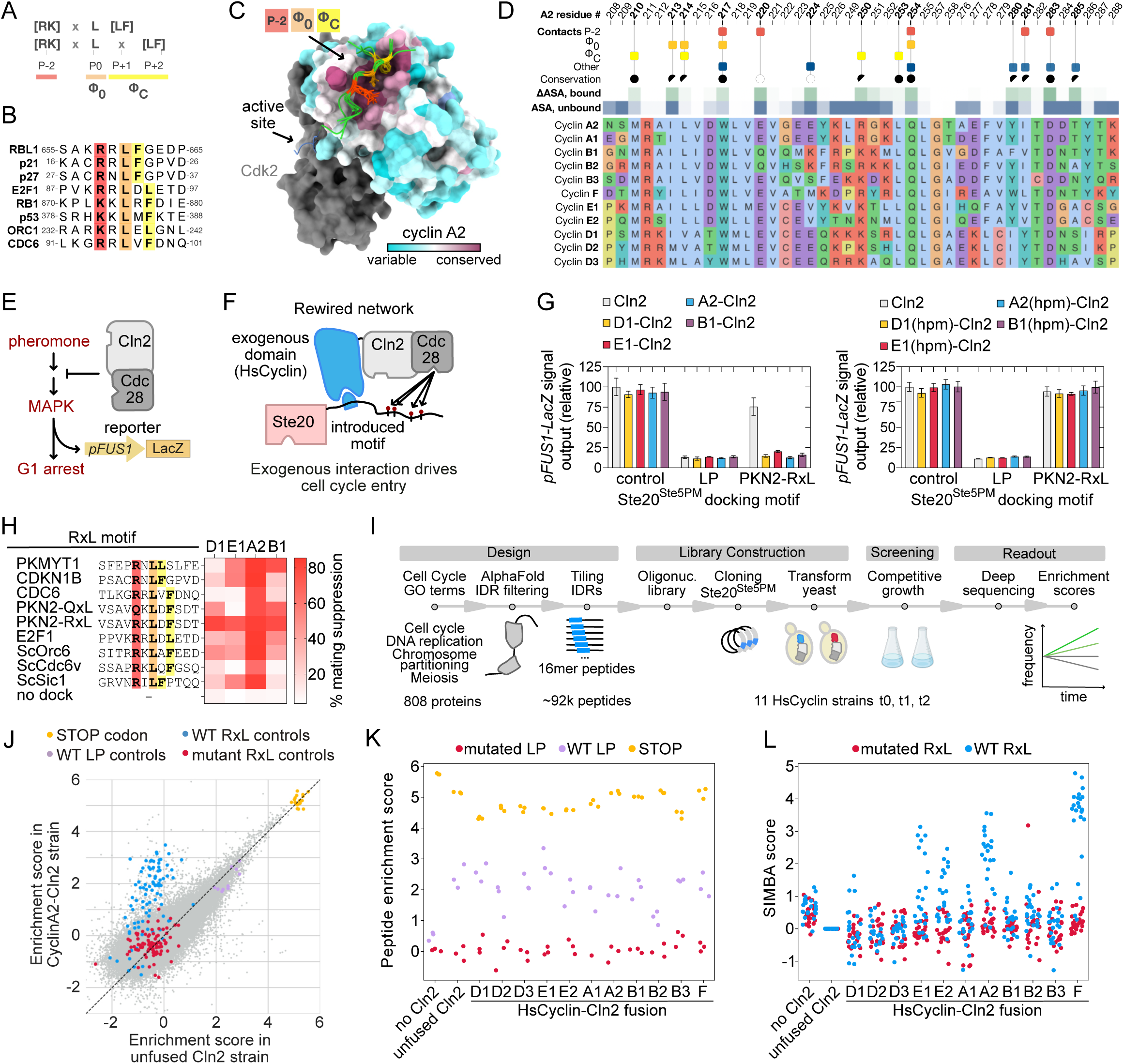
Systematic Intracellular Motif Binding Analysis (SIMBA) enables low- and high-throughput cyclin binding peptide characterization. (**A**) RxL motifs contain a central Leu at position P0 flanked by Arg/Lys at P-2 and Leu/Phe at P+1 or P+2. (**B**) Alignment of structurally characterized RxL peptides. (**C**) Superimposed structures of human cyclin A2-Cdk2 with RxL peptides bound to the hydrophobic patch (PDB: 2CCI, 6P8H, 1H24-28). (**D**) Alignment of human cyclins showing selected positions that are near the bound RxL peptides. ASA, accessible surface area. (**E**) Scheme illustrating yeast mating signaling and its inhibition by Cln2-Cdc28. (**F**) The rewired network in SIMBA, where docking between an introduced motif and a human cyclin fused to Cln2 drives Ste20^Ste5PM^ inhibitory phosphorylation. (**G**) Suppression of mating signal by wild-type and hydrophobic patch mutant (hpm) cyclins measured by a *pFUS1-lacZ* reporter. (**H**) Cyclin-motif docking measured by mating suppression relative to a Cln2 LP motif. (**I**) The cell cycle protein peptide tiling SIMBA experiment outline. (**J**) Enrichment scores of 91k peptide-encoding sequences in SIMBA competitive growth experiments with cyclin A2-Cln2 and unfused Cln2. (**K-L**) Enrichment scores of control peptides in the tiled library. Plots show raw scores (K) or Cln2-corrected enrichment scores (L).

In this study, we conduct a detailed investigation into human cyclin-binding motifs across five major cyclin families, involving over one million quantitative measurements of cyclin-motif binding strength. Using large-scale peptide binding assays, we performed unbiased screens of human proteins to identify cyclin-binding regions and their cyclin specificity. We subsequently employed saturation mutagenesis to identify their affinity and specificity determinants. Key findings were validated using fluorescence polarization (FP), cryogenic electron microscopy (cryo-EM) and in-cell phosphorylation assays. This work contributes unique insights into the rules governing substrate recruitment by human cyclin-CDK, and provides a blueprint for future studies characterizing the thousands of motif-binding domains in the human proteome.

## Results

### SIMBA enables identification of cyclin binding peptides

We applied Systematic Intracellular Motif Binding Analysis (SIMBA), a quantitative yeast-based peptide binding assay^40^, to investigate human cyclin docking interactions. In yeast (*Saccharomyces cerevisiae)*, mating and cell cycle are mutually opposing pathways, with yeast Cln2-Cdc28 phosphorylating and inactivating mating pathway proteins^41^ (**Fig 1E**). In SIMBA, this network is rewired such that an exogenous domain-peptide interaction recruits Cln2-Cdc28 to its mating pathway target, a fusion protein called Ste20^Ste5PM^ (**Fig 1F**). Consequently, mating signaling and growth arrest are suppressed in proportion to the strength of the exogenous docking interaction. These effects can be assayed by low-throughput assays (involving a transcriptional reporter, pFUS1-lacZ) or high-throughput assays (involving competitive growth)^40^. To investigate human cyclins (HsCyclins), we fused their cyclin fold domains (disabled for CDK-binding) to Cln2, and inserted their candidate docking motifs into Ste20^Ste5PM^ (**Fig 1F**). In low-throughput assays, the fusions to HsCyclins (D1, E1, A2, and B1) did not perturb Cln2 binding to its native LP docking site, but they imparted an ability to recognize a pan-cyclin binding RxL motif from PKN2 derived from Proteomic Peptide-Phage Display (ProP-PD) screening (**Fig 1G, S1A-B**). Importantly, this recognition was abolished by mutating the human cyclin hydrophobic patch. Furthermore, specificity differences among cyclins were revealed by testing multiple RxL sequences (**Fig 1H**). These results established that SIMBA could be used to study human cyclin docking.

### Design, screening and benchmarking of a tiled cell cycle protein library

To comprehensively characterize the repertoire of human cyclin binding motifs, we designed a high-throughput SIMBA experiment for screening a library of peptide sequences. We selected 808 cell-cycle-related proteins as a pool of potential cyclin binders and represented their intrinsically disordered regions (IDRs) as ∼90,000 tiled 16-mer peptides that overlap by 14 amino acids to allow motif boundaries to be delineated with 2-residue precision (**Fig 1I**). This tiled library pool, including control peptides (wild-type and mutated RxL and LP motifs, and STOP codon containing peptides), was inserted into the prey component of SIMBA, and 11 human cyclins (D1/2/3, E1/2, A1/2, B1/2/3 and F) were inserted into the bait component (**Fig 1I, S1C**). The peptide library was screened against each cyclin in competitive growth assays. Deep sequencing was performed to monitor changes in the frequency of peptides over time, from which enrichment scores were calculated. Most peptides displayed similar enrichment scores with and without a human cyclin as bait (**Fig 1J**), suggesting no bait-specific suppression of growth arrest and therefore no cyclin binding. The scores were strongly correlated across three replicate experiments (**Fig S1D**). Importantly, the control peptides showed four categories of behavior (**Fig 1J-L, S1E**). First, the variants with the highest scores in all strains were those containing STOP codons, which truncate the Ste20^Ste5PM^ protein and thereby eliminate mating signaling (**Fig 1J-K**). Second, wild-type LP motif peptides were enriched in all strains expressing Cln2 or HsCyclin-Cln2, confirming that all Cln2 fusions were expressed (**Fig 1J-K**). Third, non-functional mutant RxL or LP motifs were not enriched in any of the strains (**Fig 1J-L**). Fourth, wild-type RxL control peptides were enriched specifically in strains expressing HsCyclin-Cln2 fusion proteins (**Fig 1J**). To focus exclusively on interactions mediated by the human cyclins, in subsequent analyses we corrected the enrichment scores from the HsCyclin strains by subtracting those in the unfused Cln2 strain to obtain SIMBA scores (**Fig 1L, S1F, Table S1**). These results demonstrate that SIMBA can be scaled to study large peptide libraries.

### Cell cycle protein tiling expands the census and diversity of cyclin binding peptides

As expected, cyclin-binding peptides among the ∼90,000 peptides in the tiled library were rare, and most library peptides were not enriched in any HsCyclin-Cln2 strain (**Fig 2A, Table S1**). This is illustrated by results from RBL1, which contains three IDRs spanning 361 residues, covered by 159 peptides (**Fig 2B**). While most of these peptides were not enriched in any HsCyclin strain, two regions showed binding, with the dense peptide tiling delineating motif boundaries with 2-residue precision (**Fig 2B-C**). The first region contains a characterized RxL motif^33^, covered by five peptides that were enriched in cyclin D1, E1/2, A2, and F strains (**Fig 2B-C**). The second region, at the C terminus of RBL1, contains three peptides that were strongly enriched only in cyclin D1/2/3 strains (**Fig 2B**) confirming prior findings that this region contains a helical cyclin D-binding motif^8^. This result also demonstrates that the screen can identify peptides for distinct cyclin pockets, as cyclin D-binding helices do not bind to the hydrophobic patch^8^. Thus, combining the high-throughput SIMBA method with the tiled peptide library allows unbiased identification of binding motifs, while simultaneously defining their boundaries and cyclin specificities.

**Figure 2.**
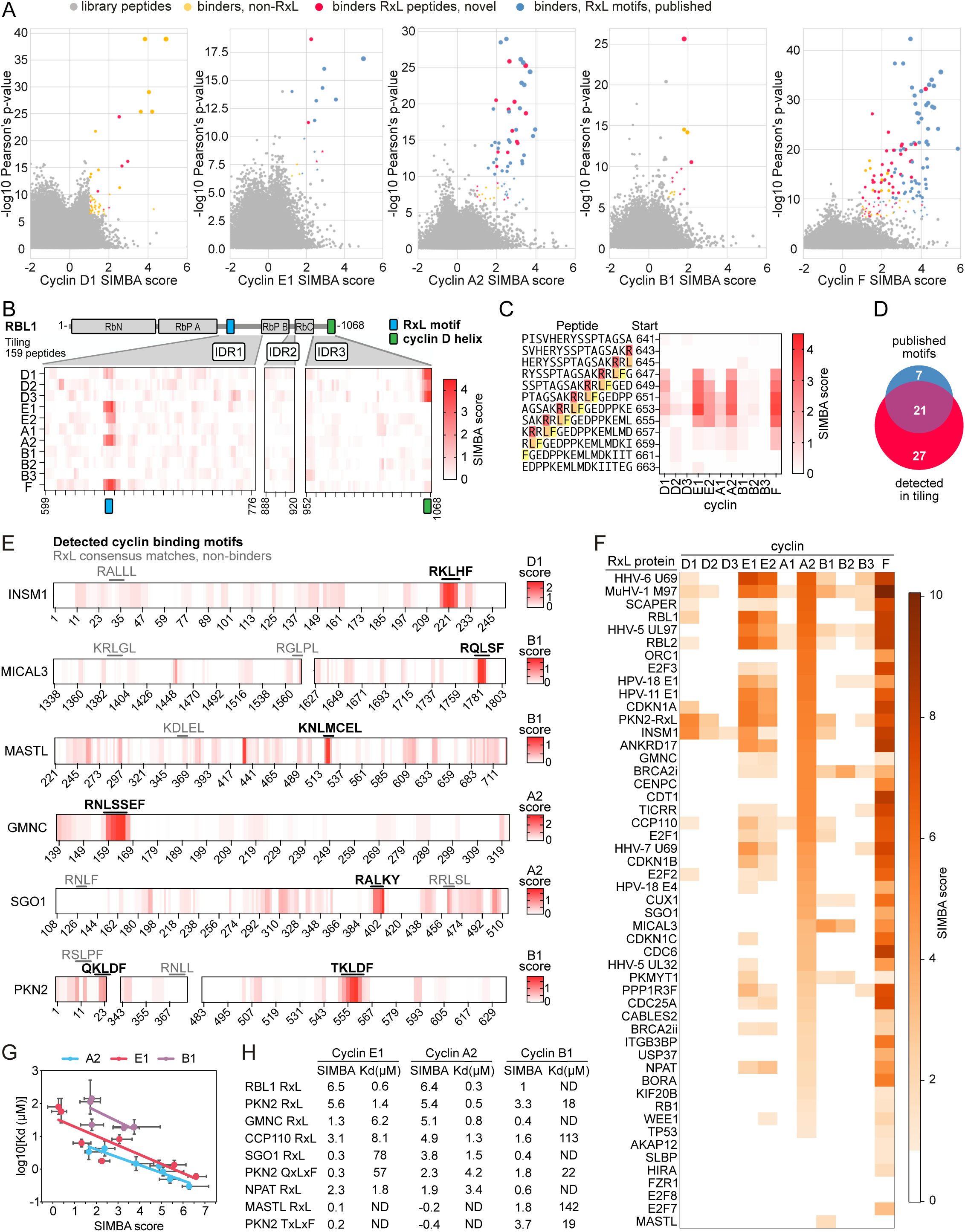
Quantitative high-throughput measurements identify cyclin binders and reveal differences in RxL binding affinity and cyclin specificity. (**A**) Identifying cyclin binders in the tiled library screen. Pearson’s p-value thresholds were 10^-10^ (high confidence; circles) and 5.5x10^-7^ (low confidence; stars). Symbols are sized by the product of the SIMBA score and the inverse log10 p-value. Control peptides are omitted. (**B**) SIMBA scores of tiled peptides from the IDRs of RBL1. Results were smoothened by plotting rolling medians of 3 consecutive peptides. (**C**) SIMBA scores (smoothened) of tiled peptides surrounding the RBL1 RxL motif. (**D**) 21 out of 28 published docking motifs and 27 novel motifs were detected as high confidence cyclin interactors. (**E**) SIMBA scores (smoothened) of tiled peptides from six proteins for the indicated cyclin. RxL motifs that bind are bold; non-binding motifs are gray. (**F**) SIMBA scores of cyclins binding to 51 RxL peptides. For clarity, only scores > 1 are shown. (**G**) Correlation between SIMBA scores (median ± 95% CI) and peptide-cyclin dissociation constants (Kd) measured by FP (mean ± 95% CI). For MASTL, the CI could not be calculated. (**H**) SIMBA scores and Kd measured by FP for multiple peptides. ND indicates that the Kd could not be determined because sufficient binding was not detected at the highest concentration (333 μM).

The screen identified 75% (21/28) of the published cyclin binding motifs in the tiled library, demonstrating its ability to capture biologically relevant interactions. In addition, 27 previously uncharacterized high-confidence motifs were discovered (**Fig 2D, Table S2**). The number of detected binding regions varied considerably between cyclins, with the highest observed for cyclins F and A2, and lower numbers for D1, E1, and B1 (**Fig 2A, Table S2**). Cyclins E1, A2, and F bound many of the published RxL motifs but D1 and B1 did not bind strongly to any. However, these cyclins did bind previously uncharacterized RxL motifs including examples of cyclin D1 binding INSM1, a reported D1 interactor^42^, and MICAL3, which bound tightly to cyclin B1 (**Fig 2E**, bold motifs). The screen also uncovered several atypical motifs. MASTL, a key mitotic regulator^43^, recognized cyclin B1 via a partial RxL consensus match that lacks a Leu or Phe at the Φ_C_ position (**<u>K</u>**N**<u>L</u>**MCEL) (**Fig 2E**). Similarly, cyclin A2 bound an **<u>R</u>**N**<u>L</u>**SSEF peptide in GMNC, a DNA replication protein^44^, and an unconventional RxLxY peptide in SGO1 (**Fig 2E**). Further, cyclin B1 bound two atypical motifs in PKN2: a QxLxF and a TxLxF, the former of which was also detected by ProP-PD (**Fig 2E, S1A-B**). Interestingly, many of these proteins also contained RxL consensus matches that did not bind any cyclin (**Fig 2E**, gray motifs). Altogether, these findings expand the census of cyclin docking motifs in the human proteome, including motif types that deviate from the typical RxL consensus, and show that a consensus match does not ensure binding.

### RxL motifs show diverse binding strengths and cyclin preferences

To directly compare different RxL motifs, we performed additional SIMBA experiments with a collection of published and newly discovered RxL motifs from human and viral proteins, screened as 13-mer peptides centered on the Φ_0_ Leu. These motifs showed diversity in both binding strength and cyclin specificity. The strongest binding peptides were generally from viral proteins, although comparable binding was shown by several human peptides such as those from SCAPER and RBL1 (**Fig 2F**). We identified three major categories of specificity for CDK-binding cyclins. First, *pan-cyclin peptides*, such as those from INSM1 and the viral protein MuHV-1 M97, which bind to members of all four families. Second, *partially selective peptides*, such as CDKN1A (p21) and RBL1 peptides that bound all cyclin types except B, or peptides from ANKRD17 and HPV-11 E1 that exhibited a more restricted preference for cyclins E and A but not D or B. Third, *cyclin-specific peptides*, which show strong preference for particular cyclin types. For example, SCAPER, ORC1, CDT1, and CENPC bound preferentially to cyclin A2, while MASTL bound only B-type cyclins. Thus, RxL motifs exhibit a variety of cyclin specificities. To validate the SIMBA findings, we performed competitive fluorescence polarization (FP) assays to measure peptide binding affinities for purified cyclin-CDK complexes *in vitro* (**Fig S1G**). We tested multiple peptides that spanned a broad range of SIMBA scores and found that these *in vivo* scores correlated well with their *in vitro* binding affinities (**Fig 2G-H**). These assays also confirmed cyclin preferences, such as cyclin B1’s exclusive ability to bind the MASTL peptide and poor binding to the RBL1 peptide (**Fig 2H**). Hence, the SIMBA data provide reliable quantification of relative binding strength and specificity, revealing novel insight into cyclin-CDK recruitment.

### Mutational scanning reveals RxL binding determinants and unexpected binding modes

To further understand peptide binding preferences of each cyclin, we conducted a comprehensive mutational interrogation of 51 peptide motifs, including published motifs and previously uncharacterized instances identified either in the SIMBA tiling screen or by ProP-PD. We designed a deep mutational scanning (DMS) analysis that included saturation mutagenesis of 22 peptides and alanine scanning of 29 others, and used SIMBA to screen a library of these mutant variants against 11 human cyclins (**Table S3-4**). As a quality check, we compared 48 control peptides present in both the tiled and mutational scanning libraries and observed high correlation between their scores in the different screens (**Fig S2A**). Pairwise comparisons of cyclins for their binding to the 7,968 distinct variants in the mutational scanning library revealed the strongest correlation between cyclins E1 and E2, followed by A2 and F, and D1 and D2 (**Fig S2B**). Notably, cyclin B3 was more similar to E1/2 than B1/2 (**Fig S2B**). Investigation of the impact of alanine mutations across the peptides confirmed the requirement for the three core positions: Arg/Lys at P-2, Leu at Φ_0_, and a second Φ_C_ hydrophobic residue at P+1 or P+2 (**Fig 3A, S2C**). Surprisingly, we found that two peptides with the sequence RxLL, a published motif from PKMYT1 (**R**N**L**LSL**F**)^45^ and one from BRCA2 found in the tiled library (**R**N**L**LNE**F**), did not depend on the Leu at P+1 and instead required the Phe at P+4 (**Fig 3A**). Furthermore, we observed the same P+4 requirement in several additional peptides from the tiled library screen, such as those from GMNC, MASTL, and CABLES2, which lacked a Leu or Phe at both P+1 and P+2 (**Fig 3A, S2C**).

**Figure 3.**
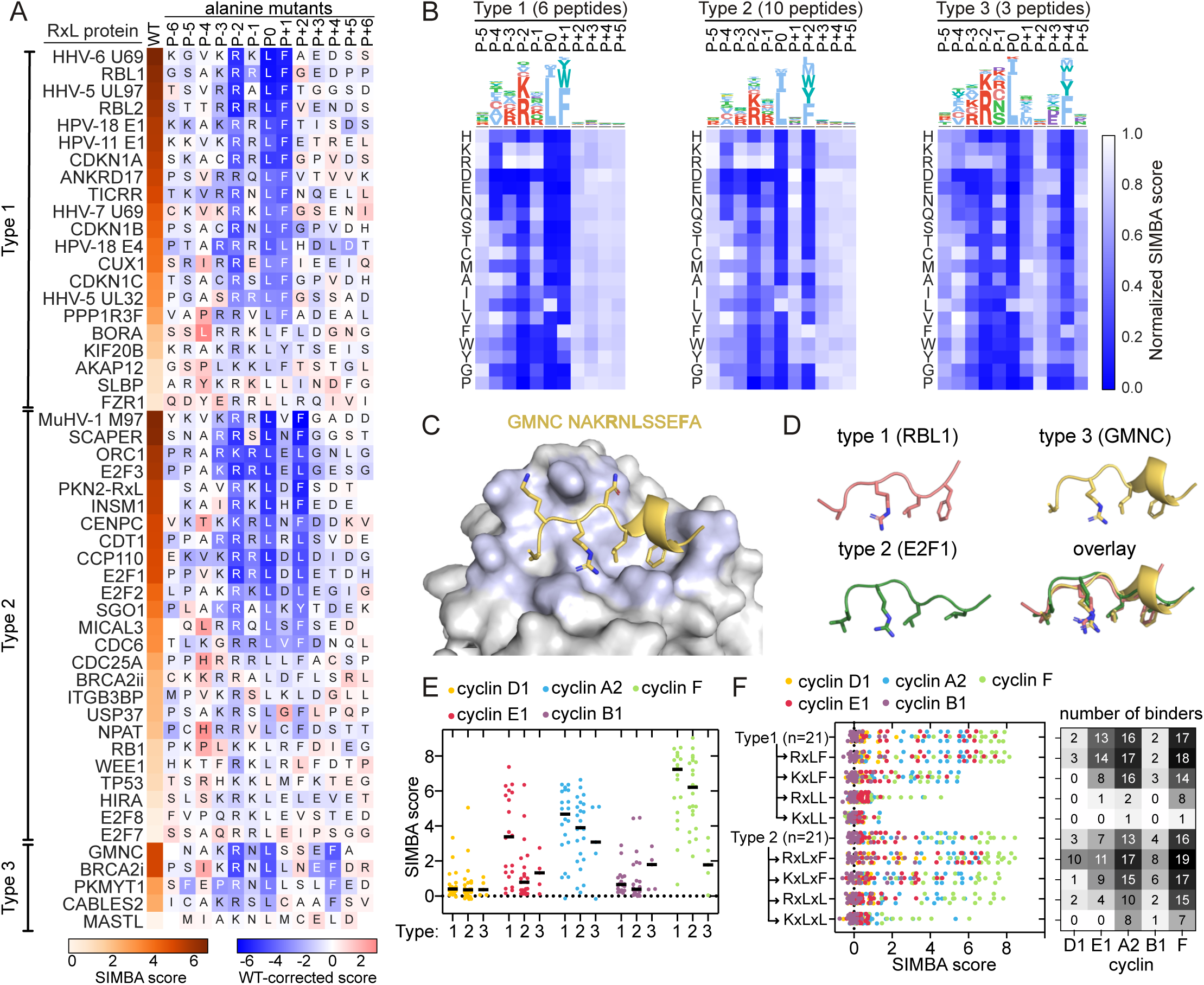
Mutational scanning reveals the impact of RxL motif core residues on cyclin binding. (**A**) RxL peptide alanine-scanning results for cyclin A2. The leftmost column shows wild-type peptide scores. Letters show the peptide sequence, and the color of the box shows the wild-type-corrected score of the peptide with Ala at that position. (**B**) Sequences logos and heatmaps showing amino acid preferences at different positions of type 1, 2, and 3 RxL motifs for cyclin A2. Data is averaged over multiple peptides subjected to DMS. (**C**) Cryo-EM structure showing the GMNC RxL peptide binding to the cyclin A2 hydrophobic patch. (**D**) Structures of cyclin-bound type 1, 2 and 3 peptides. (**E**) SIMBA scores of natural type 1, 2 and 3 peptides. Lines, median. (**F**) The impact of having either Arg or Lys at P-2 and Leu or Phe at Φ_C_ was studied by generating mutants with all combinations in the control RxL peptides.

Based on these observations, we propose grouping RxL motifs into three types — designated as type 1, type 2, or type 3 — based on whether the required Φ_C_ hydrophobic residue is at P+1, P+2, or P+4, respectively (**Fig 3A, S2C**). The averaged results of the DMS data for each type further highlight the strong preferences at the three core positions and confirm that the Φ_C_ hydrophobic residue can be provided at P+1, P+2, or P+4 (**Fig 3B, S2D, Data S1**). Interestingly, we found no example of a similar preference at P+3, suggesting that type 3 motifs bind with a specific backbone conformation. To investigate this further, we used cryo-EM to determine the structure of cyclin A2-Cdk2 bound to the type 3 peptide from GMNC. Indeed, the cyclin-bound peptide forms a single turn α-helix from P0 to P+4, which allows the Phe at P+4 to engage the same cyclin pocket contacted by the Φ_C_ hydrophobic residues in type 1 or 2 peptides (**Fig 3C-D, S3A-G**). Thus, the three RxL types allow the core position contacts to be provided by three distinct peptide conformations. The RxL type was not a strict determinant of cyclin specificity, because cyclins E1, A2, B1, and F could each bind RxL motifs of all types, although cyclin E1 showed a notable preference for type 1 motifs, and cyclin D1 did not detectably bind any type 3 motifs (**Fig 3E**). These findings uncovered a distinct motif category that was previously unrecognized.

### RxL consensus match is a poor determinant of cyclin binding

Next, we explored the contribution of the RxL consensus to cyclin binding. The tiled library contained 427 candidate motifs matching the commonly used RxL consensus sequence, ([RK]xLx{0,1}[FL]), including 287 peptides that match a stricter consensus ([^EDWNSG][^D][RK][^D]Lx{0,1}[FL]x{0,3}[EDST]) defined by the ELM resource^46,47^. We observed cyclin binding to less than 15% of the peptides with the strict consensus (**Fig S3H**), indicating that most peptides matching these consensus sequences are not cyclin binders. To further test how binding depends on the residue identity at each variable consensus position (P-2 and Φ_C_), we directly compared all four consensus combinations (RF, RL, KF, KL) in 42 parental peptides (**Fig 3F**). Both positions showed clear preferences. At P-2, Arg was preferred over Lys in both type 1 and type 2 motifs and by all cyclins, with the degree of this preference being stronger for cyclins E1 and F than for A2 and B1 (**Fig 3F**). At the Φ_C_ position, Phe was strongly preferred over Leu by all cyclins. This preference was independent of whether P-2 was Arg or Lys, but it was greater in type 1 motifs compared to type 2 (**Fig 3F**). The two positional preferences were additive, such that motifs with favored residues at both positions (RxLF, RxLxF) permitted the strongest binding and the broadest range of cyclin types. Conversely, only a minority of KxLxL motifs were binders (favoring cyclins A2 and F) and KxLL motifs were uniformly poor binders. Thus, variation in core residues alone can modulate the binding strength of RxL motifs. Conversely, peptides with optimal cores (RxLF or RxLxF) still vary in binding strength, and hence core residues alone are not sufficient to determine binding.

### Non-core positions play a key role in RxL binding strength and cyclin specificity

To obtain an overview of motif preferences for all cyclins, we grouped the DMS data by position for each of the three RxL motif types (**Fig 4A, S4A**). This analysis revealed that in addition to the three core positions, the residues at P-4, P-3, and P-1 are also strong binding determinants. Positions C-terminal to the core play a lesser role but show a preference for acidic residues that is most apparent for D and B cyclins. Compared to types 1 and 2, type 3 motifs tolerate fewer amino acids in most of the non-core positions (see numbered boxes in **Fig 4A**).

**Figure 4.**
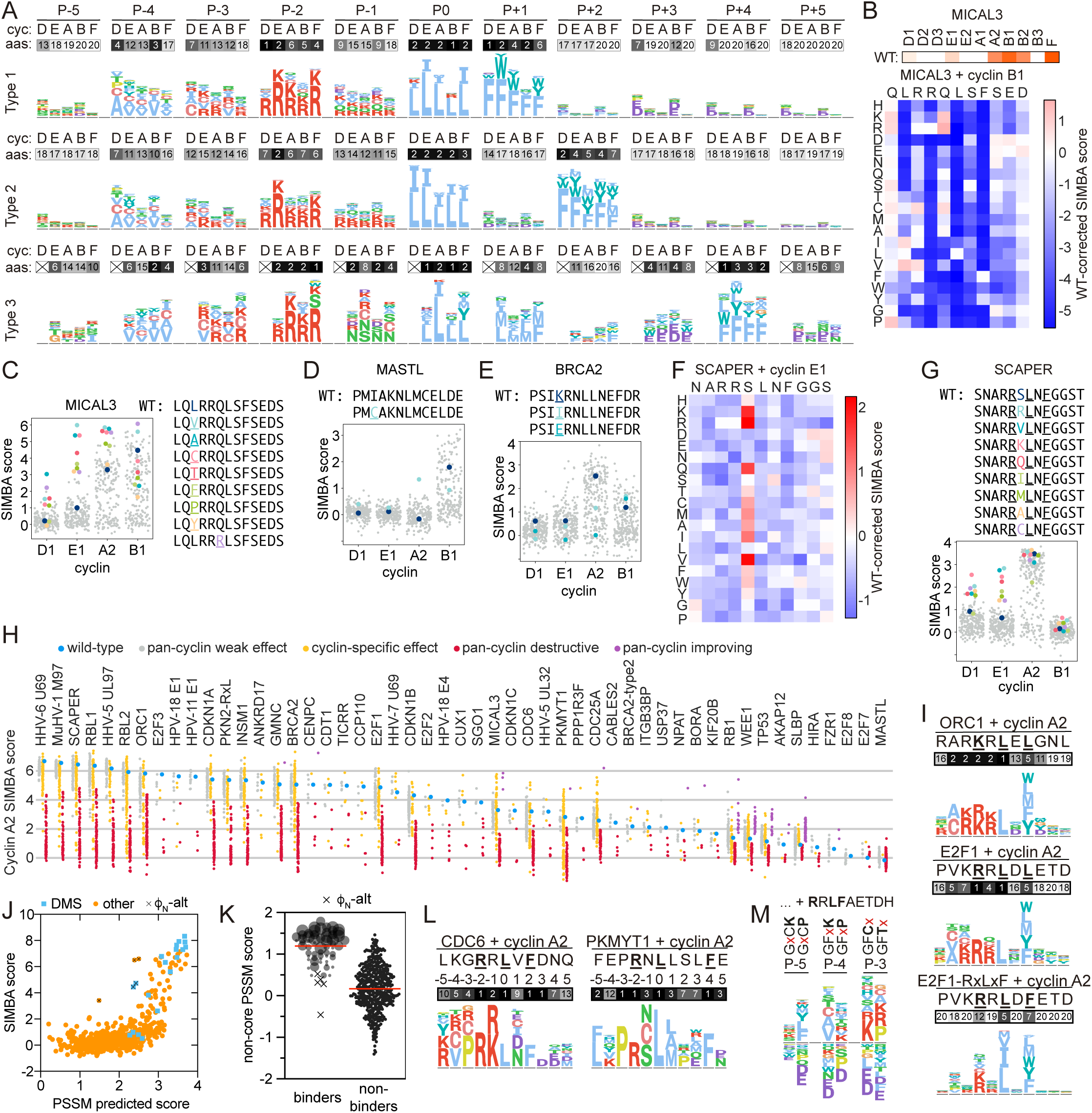
Non-core positions play a key role in determining RxL strength and cyclin specificity. (**A**) Sequence logos showing the favored residues at each position of RxL motifs for cyclins D1, E1, A2, B1 and F. DMS data of peptides where the wild-type bound the cyclin were averaged for each RxL type. Heatmaps show the average number of tolerated amino acids in each position. (**B**) SIMBA scores of wild-type MICAL3 RxL for each cyclin (top) and its DMS variants for cyclin B1 (bottom). (**C**-**E**) SIMBA scores of MICAL3 (**C**), MASTL (**D**) and BRCA2 (**E**) DMS peptides. (**F**) SIMBA scores of SCAPER peptide DMS variants binding to cyclin E1. (**G**) SIMBA scores of SCAPER peptide DMS variants. (**H**) Cyclin A2 SIMBA scores from mutational scanning of 51 peptides. Variants with scores <75% or >125% of the starting peptide were considered destructive or improving, respectively, while variants within that range were considered to have a weak effect. If this change differed by over 50% between cyclins, the effect was considered cyclin-specific. (**I**) Sequence logos representing DMS data of ORC1, E2F1 and E2F1-RxLxF motifs with cyclin A2. Heatmaps show the number of amino acids tolerated per position. (**J**) Correlation between PSSM-predicted scores and observed scores of cyclin A2 binding to wild-type RxL peptides. (**K**) Distributions of PSSM-predicted scores calculated from non-core residues only. Binder symbols are sized proportional to the SIMBA score of the parent peptide. Red lines, median. (**L**) Sequence logos representing CDC6 and PKMYT1 DMS data with cyclin A2. Heatmaps show the number of amino acids tolerated per position. (**M)** An E2F1-based RxL motif (**<u>R</u>**R**<u>LF</u>**AETDH) was appended with distinct N-terminal sequences and subjected to DMS at the position noted with ‘x’. The logos show cyclin A2 preferences at the indicated position.

At P-4, we identified strong preferences for nonpolar side chains (Ala, Val, Ile, Cys) (**Fig 4A, S4A**). Moreover, in structures of cyclin-peptide complexes, the P-4 residue is usually buried to a similar extent as the P0 Leu (**Fig S4B**), supporting a key role for P-4 as a third hydrophobic interaction which we refer to as Φ_N_. The residues favored at P-4 are largely shared by different cyclins, but there are some differences that contribute to cyclin specificity (**Fig 4A, S4C**). For example, cyclins B1 and B2 tolerate Leu and Ile at P-4 while disfavoring Ala, which is the preferred residue for cyclin D1. Notably, the cyclin B1/B2-favoring RxL from MICAL3 has Leu at P-4 (**Fig 4B**). DMS of this peptide showed that while cyclin B1 prefers hydrophobic residues Leu, Ile and Val at P-4 (**Fig 4B**), binding to cyclins E1, D1 and A2 was strongly improved by mutating the native Leu to many other residues (**Fig 4C**). Therefore, Leu at P-4 may act as a specificity filter, favoring cyclin B binding. Similarly, the cyclin B1 specific RxL motif from MASTL contains an Ile at P-4, and replacing it with Cys was sufficient to allow cyclin A2 to bind (**Fig 4D, S4D**). Suboptimal residues in the core positions of the MASTL motif might contribute to the specificity-determining effect of the P-4 residue, because cyclin A2 can tolerate Ile or Leu at P-4 in some other motif contexts (**Fig S4C**).

At P-3, most cyclins prefer Lys, Arg, Val, or Cys, although cyclin D favors small nonpolar residues (Ala, Cys, Ile) and cyclin B1 prefers Val (**Fig 4A**). The contribution of the P-3 residues to cyclin specificity is evident in data from the BRCA2 type 3 motif, where the wild-type peptide with Lys at P-3 binds both cyclin A2 and B1, but mutating P-3 to Ile or Glu increases B1 binding while significantly decreasing A2 binding (**Fig 4E**). At position P-1 (the ‘x’ of RxL), type 1 and 2 motifs prefer Arg, Lys and Gln, whereas type 3 motifs show a unique preference for Asn, Ser and Cys (**Fig 4A**), which might stabilize the adjacent helix by acting as an N-cap^48^. Despite similar preferences at P-1 for most cyclins, this position critically impacts cyclin specificity in several peptide contexts. For example, the SCAPER RxL is among the strongest cyclin A2 binders, but binds other cyclins poorly (**Fig 2F**). Interestingly, this motif gained the ability to bind cyclins D1 and E1 if the Ser at P-1 was mutated to any of several preferred residues (**Fig 4G**). Furthermore, cyclin B3 showed an unusual preference for Thr, Ser and Val at P-1 (**Fig S4A, S4E**). Thus, while having strong overlapping preferences, cyclins can differ in their tolerance for suboptimal residues, and these differences can function as specificity determinants.

Finally, to verify the impact of non-core positions, we took one peptide of each RxL type and swapped their 4-residue N-terminal flanks with those from 42 different RxL peptides, and observed that these flanks could strongly alter binding strength (**Fig S4F**). The results highlight the importance of the P-4 residue, in that cyclin D1 favored flanks with Ala at P-4 whereas cyclin B1 favored those with Val (**Fig S4F**). Most N-terminal preferences were shared by all 3 RxL types, although some flanks, such as SSLR for cyclin B1, were optimal only in the type 3 context. In analogous experiments at the C-terminal end, we observed much weaker sequence dependence, as most C-terminal flanks were tolerated (**Fig S4G**). Taken together, the DMS and flank swapping analyses demonstrate that the non-core positions contribute to binding strength and play a key part in determining the cyclin specificity of the motif.

### Core and non-core residues collectively tune RxL motif binding affinity

Next, we explored the general features controlling affinity. The mutational scanning data revealed that binding strength can be increased for all 51 peptides (**Fig 4H**), indicating that none are maximized. Overall, 51% of the mutations in RxL peptides had little to no effect, while 34% reduced binding to all cyclins (D1, E1, A2, B1 and F) and 15% produced cyclin-specific effects (**Fig 4H**). This analysis also revealed that peptides with weak core determinants had low tolerance to mutation. For instance, a KxLxL peptide from ORC1 binds strongly to cyclin A2 despite having suboptimal residues at two core positions (**Fig 2F, 3A**). DMS of this ORC1 peptide revealed limited tolerance to mutations at P-4, P-3, and P-1, in addition to the three core positions (**Fig 4I**). This suggests that the ORC1 peptide tolerates the weak core motif because it has optimal non-core residues. A similar relationship between suboptimal core residues and increased dependence on non-core positions is observed when comparing a wild-type E2F1 motif (RxLxL) to an optimized RxLxF variant (**Fig 4I**). Thus, while non-core residues are less essential than core residues, they can enhance binding strength and compensate for suboptimal cores. This concept was supported by binding determinant modeling using DMS data from each motif type (1, 2, or 3) to generate position-specific scoring matrices (PSSMs). These PSSMs allow us to sum the weighted residue preferences at all motif positions and calculate predicted scores for all natural peptides in our libraries, including additional uncharacterized peptides that contain RxL consensus matches (**Fig 4J, S4H**). We observed that, once a threshold was surpassed, there was a strong correlation between predicted and experimental scores (**Fig 4J**). We then scored these peptides using a PSSM generated from only non-core residues, and found that these non-core PSSM scores were markedly stronger for binder peptides than non-binders (**Fig 4K**). These results help explain why most RxL consensus matches are non-binders and emphasize that cyclin binding strength is a cumulative property encoded by preferences at multiple positions along the peptide motif.

### Context-specific positioning of the **Φ_N_** residue

Five outlier peptides had substantially stronger binding than predicted by their PSSM scores (**Fig 4J**, Φ_N_-alt). Interestingly, these peptides contain Pro or Gly at P-3 and a bulky hydrophobic Leu or Phe at P-5 (**Fig S4I**), which are generally disfavored (**Fig 4A**). DMS of two such motifs (CDC6 and PKMYT1) revealed a strong preference for Pro at P-3 and a greater dependence on P-5 than seen with other RxL motifs (**Fig 4L**). Co-crystal structures with E2F1 (**<u>V</u>**K**<u>R</u>**R**<u>L</u>**D**<u>L</u>**) and CDC6 (**<u>L</u>**KG**<u>R</u>**R**<u>L</u>**V**<u>F</u>**) peptides show that the P-4 Val in E2F1 and the P-5 Leu in CDC6 bind the same pocket on cyclin A2^31,33^ (**Fig S4J**). Thus, distinct binding modes could underlie the unusual N-terminal requirements. We probed the interdependence of preferences at P-5, P-4, and P-3 by DMS of these positions in parent peptides differing by a single residue. This revealed that large hydrophobic residues (Leu, Phe, Tyr) were strongly favored at P-5 when P-3 was Pro (**Fig 4M**). Testing the preferences at P-4 with or without Pro at P-3 revealed that the Pro residue reduced the usual P-4 preference for small aliphatic residues and instead imposed a weak preference for basic residues (**Fig 4M**). At P-3, Pro or Gly were favored when a Phe at P-5 was the sole hydrophobic residue but not when a Cys was available at P-4 (**Fig 4M**). Thus, the N-terminal preferences are contingent upon the available Φ_N_ residue. The flexible positioning of the Φ_N_ residue is comparable to that of the Φ_C_ residue, as both allow key pocket interactions to be encoded at multiple sites within the peptide via distinct backbone conformations, which then impose unique constraints on adjacent positions.

### Atypical RxL-like motifs lacking P-2 basic residues

While most cyclin binding peptides detected in the tiled library contained motifs conforming to the classical RxL consensus (**Fig 2A, Table S2**), several hits were only partial matches. The first category lacks a basic residue at P-2. For example, the cyclin A2-specific peptide from SAMHD1, a cyclin A2-Cdk2 substrate^34^ contains an LF sequence but has a Val at P-2 (**Fig 5A**). Despite this, the SAMHD1 peptide bound cyclin A2 with an affinity (2.68 μM Kd) comparable to many typical RxL peptides (**Fig 5B, S5A**). DMS of this SAMHD1 peptide showed that binding to cyclin A2 has an absolute requirement for the LF sequence plus distinct preferences from P-5 to P-1 (**Fig 5A**). The interaction was improved by mutating the P-2 Val to Arg or Lys, which creates an RxL consensus ([RK]QLF), yet for several cyclins binding was improved to a similar or greater degree by mutations at P-4 (**Fig 5C**). This suggests that similar atypical RxL-like motifs could also be strong interactors of other cyclins and can be tighter binders than many typical RxL motifs. Interestingly, the preferred residues at P-4 for the SAMHD1 motif are similar to those of typical RxL motifs (i.e., Cys, Ala, and Val), suggesting similar binding topologies, whereas P-5 and P-3 are more restrictive than in typical RxL motifs, perhaps to compensate for the lack of Arg/Lys at P-2 (**Fig 3B, 5A**). Using cryo-EM, we determined that the hydrophobic core (LF) of the SAMHD1 peptide contacts the cyclin A2 hydrophobic patch in a similar manner as consensus RxL motifs, and that the strongly preferred basic residues at P-5 and P-3 bind to acidic pockets distinct from the usual P-2 contacts (**Fig 5A, D, S5B-G**). We found a cyclin A2-binding SQLF peptide from CDAN1 via phage display screening that also bound exclusively to cyclin A2 in SIMBA experiments (**Fig 5E**). As with SAMHD1, alanine scanning of this CDAN1 peptide showed a key role for a P-3 basic residue (**Fig 5F**). Thus, there are multiple instances of this alternative cyclin binding mode.

**Figure 5.**
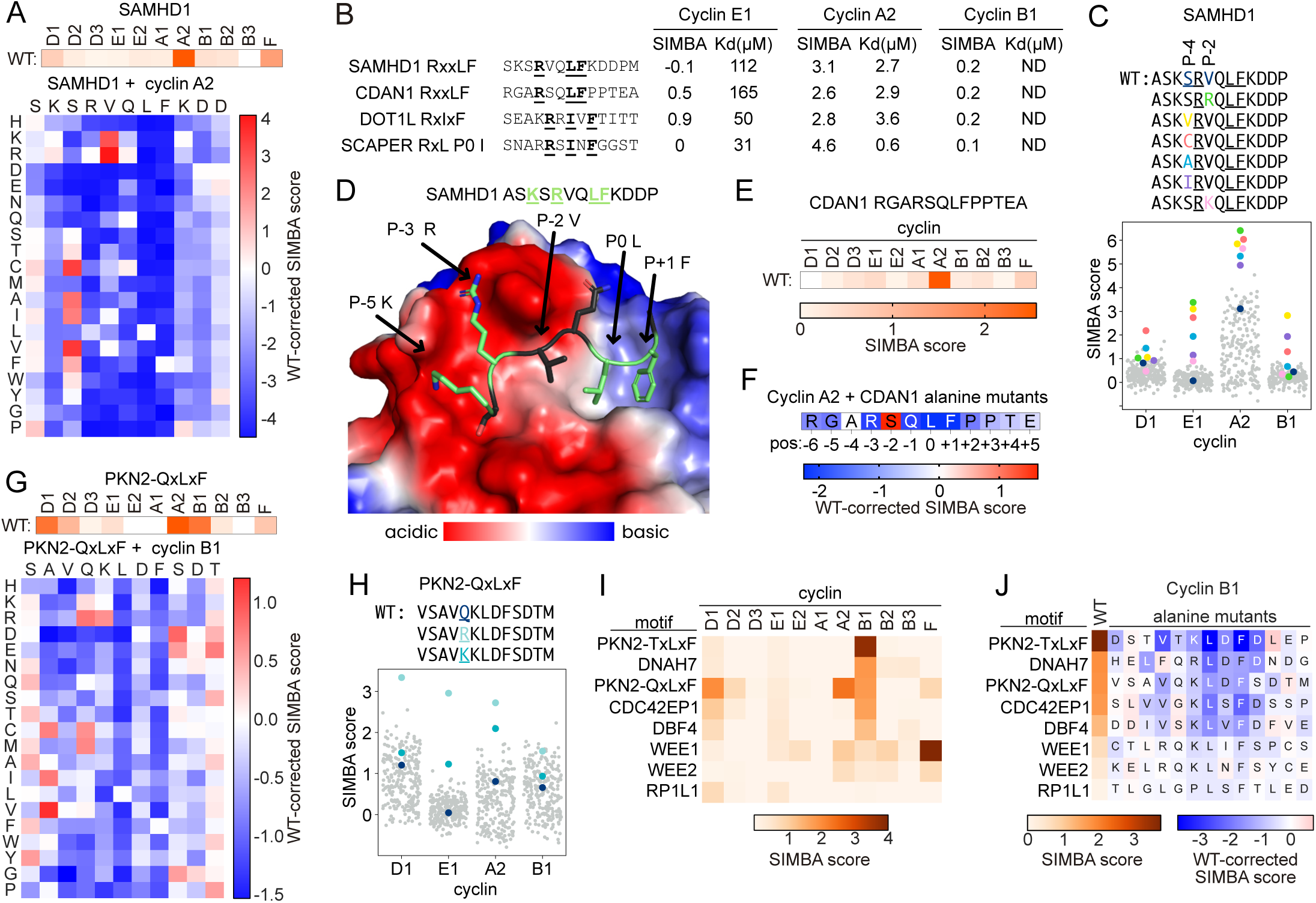
Atypical RxL-like cyclin binding motifs lacking P-2 basic residue. (**A**) SIMBA scores of SAMHD1 peptide for each cyclin (top) and the peptide DMS results for cyclin A2 (bottom). (**B**) SIMBA scores and Kd measured by FP. ND indicates that the Kd could not be determined because sufficient binding was not detected at the highest concentration (333 μM). (**C**) SIMBA scores of SAMHD1 motif DMS peptides. (**D**) Cryo-EM structure of the SAMHD1 peptide bound to cyclin A2 hydrophobic patch, colored by electrostatic potential. (**E**) SIMBA scores of CDAN1 peptide. (**F**) Alanine-scanning of the CDAN1 peptide. Cell colors represent the wild-type subtracted score of each alanine mutant with cyclin A2. (**G**) SIMBA scores of PKN2-QxLxF peptide (top) and the peptide DMS results with cyclin B1 (bottom). (**H**) SIMBA scores of PKN2-QxLxF motif DMS peptides. (**I**) The cyclin specificity of LxF-type motifs in SIMBA. (**J**) Alanine-scanning of LxF-type peptides in SIMBA with cyclin B1. Residue shading denotes wild-type-corrected scores for Ala at each position. The leftmost column shows wild-type peptide scores.

Several other atypical peptides were discovered that have an LxF motif but lack a basic residue at P-2, such as the QxLxF (VSAVQK**L**D**F**SDTM) and TxLxF (SDSTVTK**L**D**F**DL) peptides from PKN2 (**Fig S1A-B, 2E**). DMS of the PKN2 QxLxF peptide revealed that cyclin B1 binding requires the LxF residues along with strong contributions from P-4 to P-1 (**Fig 5G**). As with the SAMHD1 motif, binding of cyclin B1 to the QxLxF motif was improved by replacing P-2 with Arg to create an RxLxF motif, but Cys and Met also strengthened binding to a similar degree. Notably, cyclin E1 did not bind the native QxLxF peptide but it bound strongly to the RxLxF derivative (**Fig 5H**). Due to higher tolerance for the lack of a basic residue at P-2 by cyclins A2 and B1 than by cyclin E1, the identity of the P-2 residue could function as a specificity filter to discriminate early and late phosphorylation in the cell cycle. We investigated six additional LxF-type peptides identified in SIMBA or phage display screens and found that three of them (from CDC42EP1, DBF4, and DNAH7) bound specifically to cyclin B1 (**Fig 5I**). Alanine scanning of these peptides confirmed the essential role of the LxF sequence and revealed the importance of Val at P-3 for binding cyclin B1 (**Fig 5J**), which it also prefers in typical RxL motifs (**Fig 4A**), as well as the role of P-4 Leu in hindering cyclin A2 binding (**Fig S5H**).

### Atypical motifs with alternative residues in the hydrophobic core positions

The final category of atypical motifs, as revealed by the DMS results, showed that the core hydrophobic Φ_0_ and Φ_C_ positions can tolerate atypical residues (**Fig 3B, S3D, 4A, S4A**). Surprisingly, at Φ_C_, Trp and Tyr were generally favored over the consensus Leu (**Fig 6A, S6A**). The tiled library screen identified natural RxLY and RxLxY motifs in KIF20 and SGO1, respectively, as cyclin-binding peptides (**Fig 2E**), and alanine scanning of the SGO1 motif confirmed the Φ_C_ Tyr as part of an atypical type 2 motif (**Fig 3A**). At Φ_0_, Ile was weaker than the predominant Leu but was frequently tolerated, especially in the context of strong motifs such as SCAPER (**Fig 6B-C, S6B**). A cryo-EM structure showed that the SCAPER variant with Ile at P0 binds cyclin A2 similarly to typical RxL peptides, with the Ile residue in the pocket usually occupied by Leu (**Fig 6D**). Its binding affinity (0.61 μM Kd) was also comparable to typical RxL peptides (**Fig 2H, 5B**). While one RxI motif has been previously validated to bind cyclin F^35^, none of the natural docking motifs found previously for CDK-binding cyclins is an RxI. An *in silico* search for RxIxF motifs with optimal non-core residues identified an AK**R**R**I**V**F** peptide in DOT1L, a Cdk2 substrate^49^. This peptide bound cyclins A2 and F (**Fig 6E**), and alanine scanning confirmed the importance of the RxIxF core residues (**Fig 6F**). Thus, RxI motifs provide an additional class of potential docking sites in CDK substrates.

**Figure 6.**
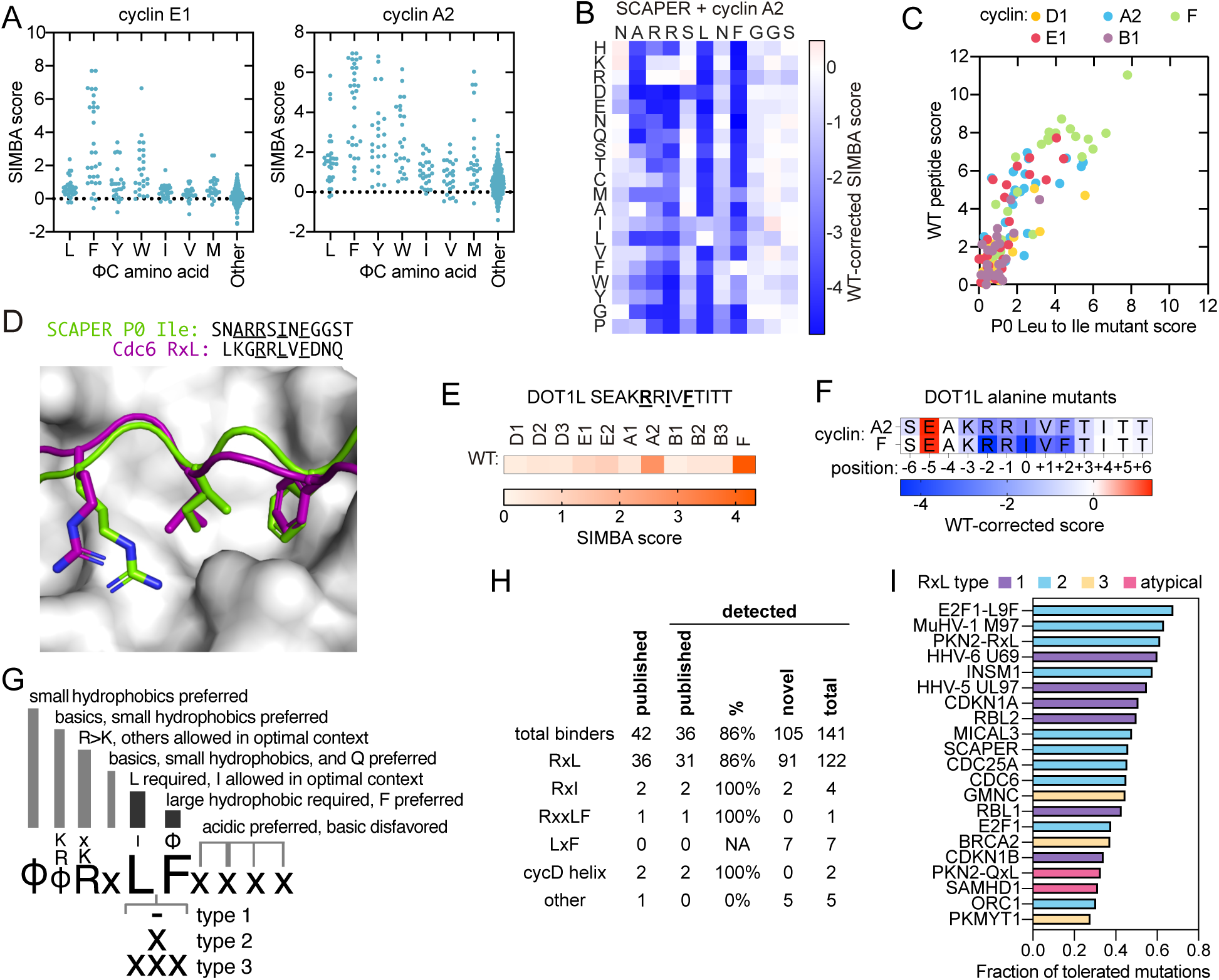
Motifs with atypical residues at core hydrophobic positions. (**A**) SIMBA scores of RxL peptides from DMS experiments with various residues at Φ_C_. (**B**) DMS results of SCAPER RxL with cyclin A2 in SIMBA. (**C**) SIMBA scores of wild-type peptides and their mutants harboring Ile at P0. (**D**) Cryo-EM structure showing SCAPER P0-Ile mutant peptide binding the cyclin A2 hydrophobic patch. (**E**) Cyclin specificity of DOT1L RxIxF peptide in SIMBA. (**F**) Alanine-scanning results of DOT1L RxIxF motif. The color of each cell reflects the wild-type-corrected SIMBA score of that position alanine mutant. (**G**) Summary of positional preferences of cyclin hydrophobic patch docking motifs. (**H**) Summary of cyclin binding motifs identified for all 11 cyclins tested by SIMBA. (**I**) The fraction of tolerated mutations, defined as mutations with SIMBA score at least 75% of the wild-type peptide, from DMS data with cyclin A2.

Collectively, these consolidated mutational scanning data indicate that the hydrophobic patch in human cyclins can recognize a greater variety of binding peptides than just typical RxL motifs (**Fig 6G**), as has also been observed with yeast cyclins^28–30^. Furthermore, *in vitro* assays showed that the atypical motifs can bind cyclins with affinities in a similar range as RxL peptides (**Fig 2H, 5B**). These findings further highlight the limitations of the previous RxL consensus, as it excludes some binding variants. Nevertheless, motifs matching the classical RxL consensus constitute the dominant fraction of cyclin binding peptides (**Fig 6H, Table S5**). Their predominance may reflect their greater sequence tolerance in non-core positions (**Fig 6I**), which could influence their likelihood of emergence during evolution.

### Cyclin docking specificity in the cell cycle

The cyclin-binding peptides display varied patterns of cyclin specificity that likely relate to their functional roles. Among the 90 natural peptides that bind the major cell cycle cyclins (D1, E1, A2, B1), the majority interacted exclusively with only a single type: 35 with cyclin A2, 10 with D1, 8 with B1, and 1 with E1 (**Fig 7A-B**). Cyclin A2 bound most of the cyclin E1 binding peptides, whereas the converse was not true, exposing a striking asymmetry in their overlap. Many peptides that exclusively bind cyclin A2 are from proteins involved in DNA replication, DNA repair, mitosis and chromosome segregation (**Fig 7B**). Among the strong pan-cyclin motifs are proteins functioning in transcription (RBL1, RBL2, INSM1) and RxL motifs from viruses (U69, M97, UL97). Cyclin D1, the first cyclin expressed in the cell cycle, binds exclusively to Rb-like helical motifs and to a small subset of RxL motifs that are mostly also recognized by cyclin E1 (**Fig 7C, S7A**). The SIMBA data suggests that the expression of cyclin E1 brings about targeting of a wider set of RxL-containing proteins. Next, cyclin A2 binds to an even wider RxL set, possibly enabling CDK-mediated phosphorylation of a broad spectrum of proteins, including those bound by cyclin E1 (**Fig 7C, S7A**). While cyclins E1 and A2 bind a wider set of RxL motifs than their predecessor, cyclin B1 interacts with a small subset of cyclin A2 RxLs (**Fig 7C-D, S7A**). Additionally, cyclin B1 exclusively binds several atypical RxL-like motifs and RxL motifs, such as the one from MASTL (**Fig S4D**).

**Figure 7.**
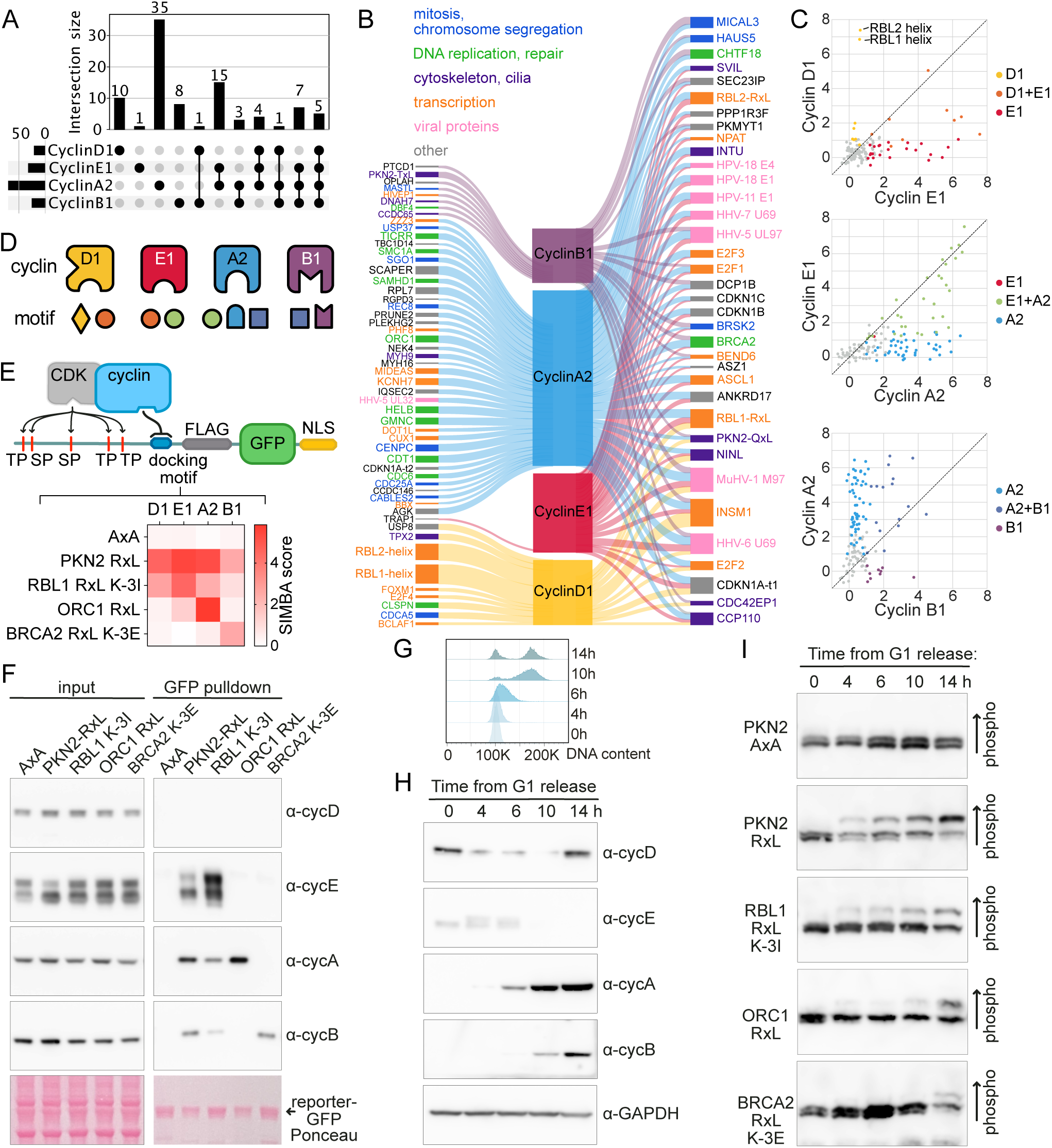
Dynamics of cyclin docking specificity in the cell cycle. (**A**) Numbers of exclusive and shared docking motifs for the major cell cycle cyclins (D1, E1, A2, B1) among wild-type peptides tested by SIMBA. (**B**) Exclusive (left) and shared (right) cyclin interactors identified by SIMBA. The width of the cyclin-substrate connecting line shows the SIMBA score. Proteins are colored by UniProt keywords. (**C**) Comparison of wild-type peptide SIMBA scores for sequential cyclins illustrates fluctuating docking specificity during the cell cycle. (**D**) Schematic illustration of cyclin docking diversification. (**E**) CDK activity reporter based on the C-terminal IDR of PKMYT1 fused to FLAG tag, GFP and nuclear localization signal (NLS). The heatmap shows SIMBA scores of the introduced docking motifs. (**F**) PKMYT1-based reporter proteins with different cyclin docking motifs were captured from human cell lysates and probed for interacting cyclins. (**G**) Cells released from palbociclib-induced G1 arrest were collected at different times and monitored for DNA content by flow cytometry. (**H**) Western blot images showing cyclin expression in synchronized cell cultures after release from palbociclib-induced G1 arrest. (**I**) The impact of docking motif specificity on phosphorylation of the PKMYT1-based reporter in palbociclib-synchronized cell cultures was monitored using Phos-tag SDS-PAGE and α-FLAG Western blot.

### Cyclin docking specificity encodes temporal outputs

To investigate how cyclin docking motif specificity influences the temporal control of CDK substrate phosphorylation during the cell cycle, we created phosphorylation reporter constructs that contain five minimal consensus CDK phosphorylation sites (from the disordered C terminus of PKMYT1), a cyclin docking motif, a FLAG tag, GFP, and a nuclear localization signal (**Fig 7E**). Pulldown experiments with these reporters confirmed that the docking motif specificities matched those observed in SIMBA experiments: (i) a mutant control motif (AxA) did not bind any cyclins; (ii) the pan-cyclin PKN2-RxL motif bound cyclins E, A and B; (iii) the RBL1 K-3I (Lys at P-3 mutated to Ile) motif bound most strongly to cyclin E; (iv) the ORC1 motif bound exclusively to cyclin A2; and (v) the BRCA2 K-3E (Lys at P-3 mutated to Glu) bound exclusively to cyclin B1 (**Fig 7F**). Next, we assayed phosphorylation of these reporters in synchronized human RPE1 cell cultures. We arrested cells in G1 with the Cdk4/6 inhibitor palbociclib, then released, and collected samples through the cell cycle^50^ (**Fig 7G**). In these experiments, cyclin D1 is detectable throughout the cell cycle, cyclin E peaks at 4 hours, cyclin A appears at 6 hours, and cyclin B appears at 10 hours (**Fig 7H**). We used Phos-tag SDS-PAGE to monitor reporter phosphorylation over time (**Fig S7B**). With the pan-cyclin RxL (PKN2-RxL), reporter phosphorylation increased gradually from 4 to 14 hours after release, whereas the control motif (AxA) caused no change (**Fig 7I**). The RBL1 K-3I RxL, which binds preferably to cyclin E1 over cyclin A2, also promoted phosphorylation from 4 to 14 hours, but not as strongly as the pan-cyclin RxL. The observation that phosphorylation of the RBL1 K-3I reporter peaks after cyclin E levels decline could be due to higher expression and activity of cyclin A2-containing CDK complexes compared to cyclin E^8^. The reporter with the cyclin A-specific ORC1 RxL showed a more abrupt increase in phosphorylation at the 14-hour time point. Finally, the cyclin B-specific RxL, BRCA2 K-3E, drove phosphorylation exclusively at the final time point and to a lesser extent than the stronger cyclin A docking motifs (**Fig 2G, 7I**). As the reporter lacking cyclin docking motifs was not phosphorylated during the cell cycle, these experiments show the importance of docking interactions in determining CDK phosphorylation and demonstrate how docking motif specificity can tune the timing of phosphorylation.

## Discussion

We have conducted a detailed investigation into the docking motif-mediated recruitment of human cyclin-CDKs, quantifying the relative binding strength of ∼100,000 peptides to 11 cyclins for a total of over one million measurements. As the largest unbiased and quantitative motif discovery and characterization screen performed to date for a protein family, it revealed insights into motif-binding domains that would not be otherwise accessible. Comprehensive screening of tiled peptides identified several new cyclin docking peptides in human proteins, including important cell cycle regulators such as MASTL, SGO1, and GMNC, showing that undiscovered motifs exist even for well-characterized motif families. Analysis of numerous cyclin-binding peptides revealed a broad diversity of binding strengths, and specificities ranging from a single cyclin to pan cyclin. In some cases, it is possible to speculate about potential physiological benefits of these differences. For example, cyclin-binding peptides from proteins that control DNA replication, such as ORC1, CDC6, and CDT1, show a strong preference for cyclin A2 over cyclin E1/2, potentially preventing premature phosphorylation before S phase. Similarly, the cyclin B1 specificity of the MASTL peptide could help ensure that CDK-antagonizing phosphatases are inactivated only in mitosis^51^. Conversely, strong, pan-cyclin binding by many viral motifs could reflect a strategy to outcompete host proteins. Variations in motif affinity might modulate the timing and/or extent of substrate phosphorylation^28,52^. All such effects likely act in concert with other factors such as the positioning and sequence context of phosphorylation sites, as well dynamic changes in localization, abundance, and activity of the cyclin-CDK holoenzymes^13,53–55^. Overall, our findings provide binding characteristics for many cyclin-binding motifs, allowing the development of new hypotheses and raising important research questions about their biological implications for the cell cycle community to explore.

Saturation mutagenesis of cyclin docking motifs in distinct peptide contexts provided an unprecedented depth of information into the specificity and affinity determinants for a motif binding pocket. The data demonstrate that numerous RxL binding modes exist with a surprisingly broad range of binding preferences, attributes, conformations, and surface contacts. Key findings include the unequal preferences at core positions, and the significant contribution of non-core positions to binding. The importance of the non-core positions is exemplified by the observation that less than 15% of peptides matching the canonical RxL consensus exhibited detectable cyclin binding. Unexpectedly, atypical binding determinants are common, including a newly-identified type 3 helical conformation and divergent motifs in which non-canonical residues replace the classical consensus at core positions. In most peptide contexts, cyclin specificity is not controlled by any single motif position; instead, subtle variations across the entire peptide collectively contribute to the binding outcome. For all tested peptides, the affinity can be increased by mutations, validating the expectation that motifs are optimized for biological output rather than for maximum binding strength. The diversity of binding determinants that modulate peptide recognition provides evolution with a broad sequence space that can be exploited to fine-tune cyclin binding from weak to strong and from highly selective to universal.

The identification of multiple binding peptides that do not conform to the consensus highlights the importance of moving beyond reliance on consensus motifs for defining binding requirements. Nevertheless, peptides matching the RxL consensus are the most frequently observed, prompting the question of why atypical variants are not more prevalent despite comparable binding affinities. Notably, they show stricter constraints at non-core positions, implying that the prevalence of the canonical consensus may be driven by the evolutionary likelihood of each binding mode. Within sequence space, there are numerous solutions for functional binding to a given motif-binding pocket, each with distinct binding attributes and evolvability. If a single binding mode within the relevant affinity range is more likely to evolve, it will dominate and define the consensus. As such, the motif consensus may represent an evolutionary bias rather than a biophysical preference.

In conclusion, our findings provide unique insights into the sequence features encoding motif binding, deepen our understanding of how CDK outputs fluctuate during the cell cycle and offer valuable guidance for developing therapeutic strategies targeting RxL motif-binding pockets^38^. The complex peptide-binding landscape uncovered here is likely to extend to other motif-binding families and our approach provides a blueprint for future studies aiming to establish general rules for motif recognition.

### Limitations of the study

Although this study provides an extensive survey of the motif-mediated interactome and binding determinants of human cyclins, some relevant interactions might still have remained undetected. First, the screens did not cover the entire proteome but instead were focused on sequences within cell cycle proteins or consensus matches in the rest of the proteome, and therefore we would not have identified atypical motifs outside of the cell cycle proteins. Second, although 75% of previously validated motifs showed binding in this study, the remaining 25% were not distinguishable from background, raising the possibility that weak but functional interactions could fall outside the detection limit of our method. Finally, our approach would not capture motifs that require post-translational modification, motifs encoded by peptides longer than 16 amino acids, or motifs that require cooperative interactions involving other peptide sequences or protein partners.

## Resource Availability

### Lead Contact

Further information and requests for resources and reagents should be directed to and will be fulfilled by Peter Pryciak and Norman Davey.

### Materials Availability

Please contact the corresponding authors for reagents. All reagents are available to other investigators.

### Data and Code Availability

Supplementary material and analysis code for data processing is available at Mendeley Data: https://data.mendeley.com/datasets/bs54ny7ns2/1. Raw counts from the deep sequencing of peptide-encoding oligonucleotide sequences are available in **Table S1** (peptide tiling SIMBA experiment) and **Table S3** (mutational scanning SIMBA experiments). The atomic models and cryo-EM maps of the and CDK2 - cyclin A2 - GMNC, CDK2 - cyclin A2 - SAMHD1 and CDK2 - cyclin A2 - SCAPER complexes have been deposited to the EM Data Resource with accession codes EMD-52201, EMD-52204, and EMD-52208 and the PDB with accession codes 9HIU, 9HIW, and 9HJ1, respectively.

## Acknowledgements

We thank Martha Cyert for critical feedback on the manuscript, Chris Richardson for support with high-performance computing, Mardo Kõivomägi for sharing cyclin-CDK expression constructs, Jonathon Pines for feedback on the manuscript and sharing cyclin constructs and RPE1-hTERT cell line, and Pau Creixell and the Creixell lab for support. N.E.D. and M.O are funded by a Cancer Research UK Senior Cancer Research Fellowship (C68484/A28159). M.O was also funded by UKRI grant EP/X042065/1 and Boehringer Ingelheim Fonds travel grant. Work by M.J.W, M.S.S, and P.M.P. was funded by grants from the NIH to P.M.P. (R01GM057769 and R01GM145795). B.J.G. was supported by a Career Development Award from the Medical Research Council of the UK, grant MR/V009354/1. Work by J.C. and C.B. was funded by the Swedish Research Council (YI: 2020-03380).

## Author contributions

M.O., P.M.P., and N.E.D. conceived and designed the work. M.O., M.J.W., M.S.S., and P.M.P. performed the SIMBA experiments. M.O. performed the FP, protein pulldown assays and phosphorylation assays. N.M.G., V.I.C. and B.J.G. performed the cryo-EM experiments. J.C. and C.B. performed the ProP-PD experiments. M.O. performed the data processing. M.O., B.J.G., Y.I., P.M.P., and N.E.D. acquired funding and supervised the experiments. M.O., P.M.P., and N.E.D. wrote the manuscript with input from all authors. All authors provided comments for the manuscript before submission.

## Declaration of Interests

The authors declare no competing interests

## Supplementary Tables legends

**Table S1.** Experimental data for the cell cycle protein peptide tiling SIMBA pooled screens.

**Table S2.** The cyclin binding motifs identified in the cell cycle protein peptide tiling SIMBA experiment.

**Table S3.** Experimental data for mutational scanning SIMBA pooled screens.

**Table S4.** Median SIMBA scores and p-values for mutational scanning SIMBA experiments.

**Table S5.** Combined set of cyclin binding motifs from peptide tiling and mutational scanning SIMBA experiments.

**Table S6.** Plasmids used in the study.

**Table S7.** Yeast strains used in the study.

**Table S8.** Cryo-EM data collection, 3D reconstruction and refinement statistics.

## Supplementary data

**Data S1.** Data from cyclin binding motifs saturation mutagenesis SIMBA experiments.

## Methods

### Recombinant protein purification

For fluorescence polarization experiments (FP) and peptide display, cyclins and Cdk2 were expressed in *E. coli* BL21 ArcticExpress DE3 cells (Agilent) and purified using the Ni-NTA purification system (Thermo Fisher Scientific). The 6xHis-tagged proteins were expressed from pET28a-based vectors (**Table S6**). For FP, Cyclins E1 and A2 were purified as Cdk2 fusion proteins^8^, while cyclin B1 and Cdk2 were purified separately and mixed subsequently. For phage display, free cyclin A2 and B1 proteins were used. The BL21 cultures were grown at 37°C to OD600 0.4, then grown in an 11°C shaker for 1h, followed by IPTG addition and overnight growth at 11°C. For cyclin B1, the cell pellets were lysed in a buffer containing 20 mM Tris-HCl pH 8.0, 800 mM NaCl, 10% glycerol, 10 mM imidazole, 0.5 mM TCEP-HCl, 1% Triton X100 supplemented with 1 mg/ml lysozyme and 1x cOmplete^TM^ protein inhibitors cocktail (Merck). Cdk2, cyclin A2-Cdk2, and cyclin E1-Cdk2 cell pellets were lysed in 50 mM Tris-HCl pH 7.4, 500 mM NaCl, 10% glycerol, 10 mM imidazole, 0.5 mM TCEP-HCl, 0.5% Triton X-100, 1 mg/ml lysozyme and 1x cOmplete^TM^ protein inhibitors cocktail (Merck) buffer. The His-tagged proteins were purified using Ni-NTA agarose and eluted with a buffer containing 200 mM imidazole. The eluates were then diluted 5x with 20 mM Tris-HCl pH 7.4, 300 mM NaCl, 0.1% Triton, 1 mM DTT buffer, concentrated using Amicon Ultra 4 10K centrifugal filters, divided into small aliquots and frozen in liquid nitrogen.

### Proteomic-peptide phage display

The purified cyclins were used as bait proteins in selections against the HD2 ProP-PD library containing ∼950,000 overlapping 16-mer peptides from the intrinsically disordered regions of the human proteome. Selections were performed following the published protocol^56^. Proteins were immobilized in 96-well plates, blocked with BSA, and incubated with the ProP-PD phage library (∼10¹¹ phages/well). After pre-clearing with GST-coated wells, the phages were transferred to cyclin-coated wells. Following washing steps, bound phages were eluted using log-phase E. coli OmniMAX, amplified with M13KO7 helper phages, and grown overnight. This selection process was repeated for three additional rounds to enrich for specific binders. The enriched phage pools were then sequenced using Illumina MiSeq, and the data were analyzed with in-house Python scripts and PepTools^56^. Peptides were ranked based on occurrence in replicate selections, sequence overlap, counts, and consensus motif identification, with further focus on medium- to high-confidence peptides meeting at least three of these criteria, particularly those containing the identified motif. We performed ProP-PD to find peptides that bind to human cyclins D1, E1, A2 and B1. Cyclin D1 and E1 failed to enrich for any specific peptides, and although several cyclin-binding peptides containing RxL motifs were identified with cyclin A2 and B1 (**Fig S1A-B**), both the A2 and B1 selections were dominated by a peptide from PKN2. Surprisingly, the PKN2 peptide (VSAV**<u>Q</u>**K**<u>L</u>**D**<u>F</u>**SDTMVQQ) contains a partial match to the hydrophobic part of an RxL motif (LxF) but lacks the basic residue.

### Systematic Intracellular Motif Binding Analysis

#### Cell cycle protein peptide tiling library design

A dataset of 808 human cell cycle proteins were identified from UniProt using keywords “Cell cycle” (KW-0131), “DNA replication” (KW-0235), “Chromosome partition” (KW-0159), and “Meiosis” (KW-0469). Proteins that lacked evidence for existence at the protein level were excluded, as were proteins with extracellular, peroxisome or mitochondrial localization. AlphaFold2 structural models^57^ were used to determine the intrinsically disordered regions of these proteins. Disordered residues were defined as those having less than 8 adjacent residues (residues within 6Å of a non-hydrogen atom of each residue). This order/disorder classification was smoothened to remove short disordered or ordered regions within longer regions with the opposite classification. Regions with the annotations “Extracellular”, “Lumenal”, “Mitochondrial intermembrane”, “Perinuclear Space”, “Signal Peptide”, “Propeptide”, “Transmembrane Region”, “Intramembrane Region”, or “Initiator Methionine” were excluded, as these are inaccessible for interaction with intracellular proteins.

The accessible intracellular disordered regions of these proteins were tiled by 16-mer peptides that overlap by 14 residues. Peptides with residues that were not covered by an adjacent overlapping peptide (e.g., peptides at the beginning or end of a disordered region) were represented by two distinct synonymous oligonucleotide sequences. The final tiled set contained 91526 peptide-encoding sequences derived from 738 unique proteins. In addition to the tiled peptides, the library incorporated 60 control peptides including 24 RxL motifs, 24 inactivated RxL mutants (RxL to AxA), 3 LP motifs, 3 inactivated LP mutants (LxxP to AxxA), and 6 sequences containing STOP codons (**Table S1**). Each of these 60 control peptides was represented by 4 synonymous oligonucleotide sequences.

#### Library design for mutational scanning

The mutational scanning was performed using 13-mer peptides containing the P0 Leu in the middle surrounded by 6 flanking N- and C-terminal residues. We performed alanine-scanning of 35 peptides and saturation mutagenesis of 25 peptides. For alanine-scanning, single alanine mutations were introduced to all positions of the 13-mer peptide. Saturation mutagenesis was carried out for either the full 13-mer or the 11-mer center region of the peptide, where variants with all 20 amino acids at each position were designed. Each peptide was encoded by three synonymous nucleotide sequences. These experiments used two distinct libraries that were tested independently. Both libraries (library 1 and library 2) included saturation mutagenesis and alanine-scanning experiments. Library 2 also included other analyses described in the text, such as comparing Arg/Lys at P-2 and Phe/Leu at Φ_C_ (**Fig 3F**), N- and C-terminal flank swapping (**Fig S4F-G**), N-terminal contingent preferences (**Fig 4M**), and scoring binding strength of previously-uncharacterized peptides that match the RxL consensus (**Fig 4J, S4H**). In addition to the experimental peptides, both libraries shared a set of 98 control peptides including 43 RxL motifs, 43 inactivated RxL mutants (RxL to AxA), 3 LP motifs, 3 inactivated LP mutants (LxxP to AxxA), and 6 sequences containing STOP codons. Each of these 98 control peptides was represented by 3 synonymous nucleotide sequences.

#### Cloning and yeast strains

The yeast strains used in this study are in W303 background and are listed in **Table S7**. Standard methods were used for growth and genetic engineering of yeast. Yeast were grown at 30°C in yeast extract/peptone medium with 2% glucose (YPD) or in synthetic complete medium (SC) lacking uracil (or uracil and histidine) supplemented with 2% glucose or raffinose. The human cyclin reading frames were cloned between GST and yeast *CLN2* in the *P_GAL1_-GST-CLN2* expression cassette (pPP4343). The intrinsically disordered N- and/or C-termini of the human cyclins were truncated to remove localization and degradation signals. To prevent the human cyclins from interacting with yeast Cdk1, a triple alanine mutation was introduced in the conserved CDK-binding interface of each cyclin (at positions corresponding to cyclin A2 residues Lys266, Glu295, and Phe304), based on prior mutations in yeast cyclins^20,58^; an exception was cyclin F, which lacks these conserved residues and does not bind CDKs. The cyclin expression constructs were harbored on replicating (CEN/ARS) plasmids or were integrated at the *HIS3* locus in yeast. The cyclin docking peptides were inserted into Ste20^Ste5PM^ chimera and expressed in yeast from a low copy number plasmid with *URA3* selection. 13-mer or 16-mer peptide sequences for mutational scanning or cell cycle protein peptide tiling libraries, respectively, were flanked by GSGG flexible linker sequences in the Ste20^Ste5PM^ construct. Oligonucleotide pools encoding docking peptides were ordered from GenScript and were amplified by PCR (10 cycles) using primers that anneal to the common flanking linkers and include MluI and SphI restriction sites. The PCR products were digested with MluI and SphI, treated with calf intestinal phosphatase, and then ligated into the MluI/SphI-digested Ste20^Ste5PM^ expression vector (pPP4745). The ligation products were transformed into E. coli (XL-10 Gold Ultracompetent Cells; Agilent/Agilent Technologies) and plated on LB+Amp plates. To obtain a good representation of the designed library, at least 30-fold greater number of bacterial transformants compared to the number of variants in the library were harvested and used for plasmid preparation from the pool of colonies. The plasmids used in this study are listed in **Table S6**.

#### β-galactosidase expression assays

As a low-throughput quantitative readout of cyclin binding to the Ste20^Ste5PM^ protein, mating pathway signaling was measured in yeast strains that contain a *FUS1-lacZ* transcriptional reporter construct driving β-galactosidase expression^59^. For this, yeast cultures were grown in SC medium containing raffinose and, at OD_600_ 0.7, the exogenous cyclin expression was induced by adding galactose (2% final). After 90 minutes, mating signaling was activated by adding α-factor to 50 nM and the cultures were grown for an additional 45 minutes. Then, the yeast cells were pelleted and resuspended in 60 mM disodium phosphate, 40 mM monosodium phosphate, 10 mM KCl and 1 mM magnesium sulfate buffer. The cells were permeabilized by addition of chloroform (to 10%) and sodium dodecyl sulfate (to 0.008%), and then ortho-nitrophenyl-β-galactoside was added (to 0.9 mg/ml) as a spectrophotometric substrate for β-galactosidase activity. The reactions were incubated at 30°C. β-galactosidase activity was estimated by measuring formation of ortho-nitrophenol in the reaction supernatant at OD_420_.

#### Yeast competitive growth experiments

The pooled Ste20^Ste5PM^-peptide expression plasmid libraries were transformed into yeast strains harboring integrated expression cassettes for different HsCyclin-Cln2 fusions. The transformation mixture was plated on SC medium plates lacking uracil, and grown at 30°C for three days. A dilution series from the transformation was used for colony counting to ensure that the number of yeast transformants exceeds the number of peptides in the library at least 10-fold. The transformed cells were collected from the plates, diluted to 50 ml SC-URA containing raffinose and grown at 30°C for 4-8 hours. Then, the 50 ml cultures were diluted to OD_660_ 0.05 and grown overnight. At OD 0.8-1, HsCyclin-Cln2 expression was induced by adding 2% galactose. After 75 minutes, 40 ml of the culture (∼ 3 × 10^8^ cells) was collected by centrifugation and flash frozen in liquid nitrogen to obtain the t0 sample. At the same time, α-factor at 500 nM final concentration was added to the culture to begin the competitive growth experiment. The cultures were grown at 30°C and diluted every 12 hours to keep the OD_660_ below 1. Aliquots (20 ml; ∼ 3 × 10^8^ cells) were harvested at 20 and 32 h from α-factor addition. The harvested cells were centrifuged, washed with water, moved to 1.5 ml tubes, where the cells were pelleted by centrifugation, the cell pellets were frozen and stored at -80°C.

#### Sample preparation for deep sequencing

DNA was purified from yeast cells using the Zymo Research ZR Plasmid Miniprep Kit (#D4015). Frozen cell pellets were thawed and suspended in 200 μl of solution P1, and then were lysed using Zymolyase (0.2 units/μl; Zymo Research #E1005) for 1.5 h at 37°C, before proceeding with the remaining purification steps. Samples of plasmid DNA (4 μL) were subjected to PCR (17 cycles, 50 μL total volume) with primers that include standard P5 and P7 sequences for binding to Illumina flow cells during next generation sequencing. The forward primer included a P5 sequence followed by an Illumina sequencing primer binding site, a 6-nucleotide bar code, and an upstream plasmid-annealing sequence; the reverse primer included a P7 sequence followed by a 6-nucleotide i7-index sequence, an i7 sequencing primer binding site, and a downstream plasmid-annealing sequence.

Aliquots (5 μl) of the PCR products were run in 1.2% agarose gels to confirm the presence of the desired product, and the remainders were purified using Zymo Spin I columns (Zymo Research #C1003-250) and eluted in 10 mM Tris-HCl, pH 8. The concentration of the eluted products was measured using nanodrop, and a mixture was prepared containing equal amounts of each DNA product. The PCR amplicon libraries were sequenced at Novogene.

#### Sequencing data analysis

2FAST2Q was used to obtain the counts of different peptide-encoding oligonucleotide sequences from the deep sequencing data^60^. To calculate enrichment scores, custom Python scripts were used for the following data processing. (1) Oligonucleotides with <40 reads at t0 were dropped from further analysis. (2) The frequency of each oligonucleotide within a sample at t0, t20 and t32 was calculated. (3) The log2 fold change of t20 and t32 frequencies from the t0 frequency was calculated. (4) The log2 fold change values were z-normalized within each sample. For mutational scanning library experiments, z-normalization used the mean and standard deviation of the P0 Leu mutwants; for tiled library experiments, z-normalization used the mean and standard deviation of the entire oligonucleotide population. (5) A quadratic regression (specifically, a degree 2 polynomial fit performed using numpy.polyfit) was used to correct for nonlinearity between z-scores in HsCyclin-Cln2 strains versus the unfused Cln2 strain. These corrected values are called enrichment scores. For mutational scanning libraries, this regression was performed using data from mutant RxL and LP peptides, wild-type LP peptides and STOP codon containing variants; for the tiled library, the full data set was used. (6) For each oligonucleotide, the score in the unfused Cln2 strain was subtracted from the HsCyclin-Cln2 strain score to obtain the final SIMBA score. The z-normalization and nonlinearity corrections were performed differently for data from the peptide tiling and mutational scanning experiments because in the tiling library the vast majority of variants are non-binders and thus are comparable between unfused Cln2 and HsCyclin-Cln2 strains (**Fig 1J**), whereas the mutational scanning libraries contain a large fraction of HsCyclin binding peptides. Therefore, the z-normalization and nonlinearity correction of the mutational scanning experiment data was performed using the control set variants that are expected to score identically in unfused Cln2 and HsCyclin-Cln2 strains (wild-type LP, mutant LP and mutant RxL peptides, and STOP codon containing variants). As described above, we created two distinct mutational scanning libraries that were tested in independent experiments. To compare SIMBA scores for peptides in the different libraries, we used all peptides shared between the two libraries (199 total) to determine correction factors that were used to adjust the scores from library 1 to fit those in library 2. The equations used for these corrections are provided in **Table S4**.

In mutational scanning experiments, each peptide is represented by 3 synonymous oligonucleotides, and the plotted data represent the median score for these 3 synonyms from two replicate experiments. Independent t-tests were performed using the set of 6 scores (3 synonyms in 2 replicates) to test if the median peptide score is greater in a given HsCyclin-Cln2 strain than in the unfused Cln2 and no cyclin strains. In the tiled library screens, each peptide is encoded by a single oligonucleotide, but adjacent 16-mer peptides overlap by 14 residues. With this library, the statistical tests were performed on sets of three consecutive peptides that group a given peptide with its preceding and subsequent peptides; these statistical tests were not performed for cyclin strains that were tested in only one replicate with the tiled library (A1, B2, B3, E2, D2, D3). The plotted data (**Fig 2B, 2C, 2E**) show smoothened results that represent the rolling medians of each set of 3 consecutive peptides. The p-values from the two t-tests (i.e., HsCyclin vs. unfused Cln2 and HsCyclin vs. no cyclin) were combined using Pearson’s method. Binders were defined as those with a SIMBA score >1 and Pearson’s p-value <6.36x10^-6^ or <5.52x10^-7^ for mutational scanning and tiled library screening, respectively (these correspond to p <0.05 after Bonferroni correction for library size). In addition, binder peptides from the tiled library screen that had a Pearson’s p-value <1x10^-10^ were designated as “high confidence” binders. While the tiled library included an additional set of (untiled) RxL control peptides, only the tiled peptides were included in **Fig 2A** and **Table S2** as identified cyclin binders. Finally, to obtain a consolidated list of cyclin binding motifs detected in the tiled library screen, overlapping binding peptides were collapsed to the minimal shared sequence. Then, the SIMBA score and Pearson’s p-value of the shared motif were calculated as the median of values from the individual overlapping binding peptides.

To create logos of sequence preferences, SIMBA scores were transformed into a preference PSSM as described previously^52^. First, the score for each amino acid variant was normalized to the lowest and highest scores in a given motif array. Then, these normalized scores were converted to a frequency metric by dividing each by the sum of all scores at the same position. Finally, the frequency metric was converted to a preference metric by subtracting 0.05, so that a neutral preference is represented by zero, favored residues are positive, and disfavored residues are negative. These preference scores were used to generate sequence logos via a web-based tool (http://slim.icr.ac.uk/visualisation/index.html). To compare predicted versus observed binding strength, SIMBA scores were normalized as described above and then transformed into a difference PSSM^52^ by subtracting the average normalized score of all residues at a given position from the value of each residue at that position: (difference score) = (residue score) – (position average). Then, to obtain a prediction for a given peptide, the corresponding PSSM values for each residue at each position was summed across all motif positions to calculate the predicted score (PSSM sum).

### Fluorescence polarization

Unlabeled peptides and N-terminally FITC-Ahx labeled PKN2-RxL peptide with >95% purity were ordered from GenScript. Fluorescence polarization was measured in a buffer containing 25 mM Hepes-KOH pH 7.4, 150 mM NaCl, 5 mM DTT and 0.05% CHAPS. The assays were carried out in black, low volume, non-binding, 384-well plates (4514, Corning). After mixing, the reactions were incubated at 37°C in the dark for 30 minutes before measuring fluorescence polarization using PHERAstar FSX microplate reader. Peptide binding affinities were measured by dosing unlabeled peptides that compete with the FITC-PKN2-RxL peptide for binding to the cyclin hydrophobic patch. The optimal concentration of cyclin proteins in the experiment that allows sufficient resolution in fluorescence polarization was determined by measuring the binding affinity of the cyclins to FITC-PKN2-RxL peptide. In the competitive binding experiments, recombinant cyclin B1 was mixed at equimolar ratio with Cdk2 and used at 4 µM concentration, cyclin A2-Cdk2 was used at 0.5 µM, and cyclin E1-Cdk2 at 0.15 µM. The fluorescence polarization experiments were performed in at least two replicates. The data was analyzed and plotted using Prism (Graphpad).

### Generating CDK reporter expression cell lines

CDK activity reporters based on the intrinsically disordered C terminus of PKMYT1, where the native RxL motif^45^ was replaced with different cyclin docking motifs. The CDK reporters were fused to a 3x FLAG tag, EGFP, and a nuclear localization signal, and expressed from a doxycycline-inducible lentiviral vector (pCW57.1). The virus was produced as follows: 7 x 10^5^ HEK293T/17 cells were seeded to 6-well plate and in 24 hours were transfected using lipofectamine 3000 (Thermo Fisher Scientific) with psPax2 and pMD2G as packaging plasmids. 6 hours after the transfection, the medium was replaced with 3 ml fresh DMEM. The medium containing the virus was collected 2 days after transfection and was used to transduce RPE1-FRT cells.

RPE1-FRT cells were cultured in high glucose DMEM supplemented with 110 mg/l pyruvate, 3.7 g/l sodium bicarbonate, and 10% fetal bovine serum in a 37°C incubator with 5% CO_2_. 24 hours before transduction, 2 x 10^5^ RPE1-FRT cells were seeded to a 6-well plate. For transduction, the medium of RPE1-FRT cells was replaced with the lentiviral supernatant and supplemented with 8 µg/ml polybrene, followed by culturing the cells for 2 days. Then, the transduced RPE1-FRT cells were collected by trypsinizing and seeded to a 6 cm dish in DMEM supplemented with 1 µg/ml puromycin. Prior to seeding for experiments, the transduced RPE1-FRT cells were split at least 3 times and cultured in the presence of puromycin to allow selection of the transduced cells.

### RPE1 cell cycle synchronization

For cell cycle synchronization, 2.5 x 10^5^ RPE1-FRT cells containing the CDK reporter expression cassettes were seeded to 10 cm dishes and grown for 32 hours. Then, the medium was replaced with DMEM containing 150 nM palbociclib (Selleck Chem) to arrest cells in G1 phase and 1 µg/ml doxycycline to induce the expression of CDK reporters. The cultures were incubated at 37°C for 24 hours, after which the palbociclib-containing medium was removed, the cells were washed 3 times with 10 ml DMEM and the cells were grown in DMEM supplemented with 1 µg/ml doxycycline at 37°C. At 0, 4, 6, 10 and 14 hours, the cells were collected by trypsinization, washed with Tris buffered saline and flash frozen for immunoblotting. For propidium iodide staining, the cells were fixed with 70% ethanol and stored at -20°C.

### Propidium iodide staining and flow cytometry

The ethanol-fixed cells were washed twice with PBS and resuspended in 300 µl solution containing 5 µg/ml propidium iodide and 25 µg/ml RNase A in PBS. The mixture was incubated at 37°C in the dark for 30 minutes and analyzed by flow cytometry using BD LSR Fortessa. Data from at least 10 000 cells was collected and analyzed using FlowJo.

### Co-immunoprecipitation

For CDK reporter co-immunoprecipitation experiments, the RPE1-FRT-based cell lines were seeded to 15 cm dishes in DMEM, at 40% confluency, 1 µg/ml doxycycline was added to the medium to induce the reporter expression and the cells were collected and frozen at 80% confluency 24 hours after induction.

The cells were lysed in buffer containing 10 mM Tris-HCl pH 7.4, 150 mM NaCl, 0.1% NP-40, 5 mM EDTA, 1 mM DTT and cOmplete^TM^ protein inhibitors cocktail (Merck). The total protein concentration in the lysate was measured using Pierce™ Bradford Plus Protein Assay Reagent (Thermo Fisher Scientific). The lysates were diluted to 2 mg/ml total protein. For co-immunoprecipitation, 500 µg of cell lysate was mixed with 10 µl ChromoTek GFP-Trap® Magnetic Agarose beads (Proteintech) that had been equilibrated with the lysis buffer. The mixture was incubated on a rotator at 4°C for 1 hour. Then, the GFP-trap magnetic beads were collected using a magnet, the lysate was removed and the beads were washed three times with 500 µl lysis buffer. The immunoprecipitated proteins were eluted using Laemmli SDS-PAGE sample buffer and were subjected to SDS-PAGE and immunoblotting.

### Immunoblotting

To analyze the phosphorylation of the CDK reporters in the cell cycle, the synchronized RPE1-FRT cells expressing the reporter-GFP fusion proteins were lysed in a buffer containing 10 mM Tris-HCl pH 7.4, 150 mM NaCl, 1% Triton X100, phosSTOP phosphatase inhibitor cocktail (Roche) and cOmplete EDTA-free protease inhibitor cocktail (Merck). The lysate was cleared by centrifugation and the total protein concentration was measured using Pierce™ Bradford Plus Protein Assay Reagent. For immunoblotting, 20 µg of cell lysate was resolved using Phos-tag SDS-PAGE with gels containing 8% acrylamide, 25 µM Phos-tag (Alpha Laboratories) and 50 µM MnCl_2_^61^. Following electrophoresis, the gels were washed twice with 10 mM EDTA. The proteins were transferred to nitrocellulose membrane using iBlot 2 (Thermo Fisher Scientific). After transfer, the membrane was blocked using 5% fat-free milk solution in TBS-T, followed by overnight incubation at 4°C with the primary antibody solutions. The following primary antibodies were used: sc-20044 (Santa Cruz Biotechnologies) at 1:500 for Cyclin D1, sc-247 (Santa Cruz Biotechnologies) at 1:500 for cyclin E, sc-271682 (Santa Cruz Biotechnologies) at 1:500 for cyclin A, sc-245 (Santa Cruz Biotechnologies) at 1:500 for cyclin B, anti-GAPDH (D16H11) XP® Rabbit mAb #5174 (Cell Signaling Technologies) at 1:1000 for GAPDH, a-FLAG tag (D6W5B) Rabbit mAb (Cell Signaling Technologies) at 1:1000. Then, the membrane was washed 5 times with TBS-T, incubated with secondary antibody solutions for 1 hour, and washed again 5 times with TBS-T. HRP-conjugated anti-mouse IgG (#7076, Cell Signaling Technologies) at 1:2000 and HRP-conjugated anti-rabbit IgG (#7074, Cell Signaling Technologies) at 1:10000 were used as secondary antibodies. The antibodies were detected using SuperSignal™ West Pico PLUS Pico Chemiluminescent substrate (Thermo Fisher Scientific).

To validate the protein bands observed in Phos-tag SDS-PAGE, lysates from asynchronous RPE1-FRT cultures expressing the CDK reporter proteins were dephosphorylated using lambda protein phosphatase. For this, 20 µg of the lysates prepared without phosphatase inhibitors were supplemented with 1 mM MnCl_2_, 1x NEB PMP buffer and 8 U/µl lambda protein phosphatase (New England Biolabs). The reactions were incubated at 30°C for 30 minutes and stopped by addition of Laemmli SDS-PAGE sample buffer.

### CDK2-cyclin A expression and purification for cryo-electron microscopy

Cdk2 and cyclin A were expressed separately and combined later during purification. The human Cdk2 construct (residues 1-282) was provided by John Chodera, Nicholas Levinson and Markus Seeliger (Addgene plasmid #79726; http://n2t.net/addgene:79726; RRID: Addgene_79726)^62^. Human cyclin A2 containing an N-terminal His_6_ tag was cloned into a vector suitable for bacterial expression. For each construct, 10 ml of LB supplemented with 100 µg/ml ampicillin and 34 µg/ml chloramphenicol were inoculated with a single colony of transformed *E. coli* Rosetta (DE3) pLysS cells and incubated at 37°C overnight. The whole overnight culture was used to inoculate 3 l of LB supplemented with 100 µg/ml ampicillin and 34 µg/ml chloramphenicol that were incubated at 37°C. Cultures were induced at OD ∼ 0.6 with 1 mM IPTG and incubated at 24°C overnight for protein expression. Cultures were harvested by centrifugation at 10,000 x g for 10 min at 4°C. Pellets were snap frozen in liquid nitrogen and stored at -80°C.

To form the cyclin A-Cdk2 complex, 7.8 g of Cdk2 pellet and 9.5 g of cyclin A pellet were combined and resuspended in purification buffer (50 mM HEPES pH 7.5, 180 mM NaCl, 5% (v/v) glycerol, 2 mM MgCl_2_, 2 mM DTT) supplemented with protease inhibitors and DNAseI. Cells were then lysed by sonication and the lysate was clarified by centrifugation at 18,000 rpm for 30 minutes at 4°C, before spinning the supernatant for a further 15 minutes. The clarified lysate was supplemented with 10 mM imidazole and then loaded onto an equilibrated 5 ml HisTrap column (Cytiva). The column was washed with purification buffer supplemented with 20 mM imidazole. Complexes were eluted using a linear gradient of 20-300 mM imidazole across 60 ml, then the column was washed with purification buffer supplemented with 300 mM imidazole for a further ∼30 ml. Fractions containing cyclin A-Cdk2 complexes were combined and incubated overnight with TEV protease to cleave the tags. After cleavage, complexes were buffer exchanged to gel filtration buffer (20 mM HEPES pH 7.5, 150 mM NaCl, 5 % v/v glycerol, 2 mM MgCl_2_, 2 mM DTT), centrifuged at 4,000 rpm for 15 mins at 4°C, and then incubated with equilibrated Ni-NTA Superflow beads (Qiagen) at room temperature for 30 min to remove His_6_-tagged TEV protease. The flow-through was then concentrated, flash frozen in liquid nitrogen and stored at -80°C. Complexes were thaw and further purified by gel-filtration using a Superdex 200 Increase 10/300 column (Cytiva). Cyclin A-Cdk2-containing fractions with a concentration of 2 mg/ml (1 Abs = 1 mg/mL) were aliquoted, snap frozen in liquid nitrogen and stored at -80°C.

### Cryo-EM grid preparation

Cryo-EM samples were prepared on UltrAuFoil 1.2/1.3 300 mesh holey gold grids (Quantifoil Microtools). Samples were prepared by mixing the cyclin A-Cdk2 complex (5x dilution from 2 mg/mL stock) with 300 μM of RxL peptide in cryo-EM buffer (20 mM HEPES-NaOH pH 7.5, 150 mM NaCl, 2 mM MgCl_2_). 4 μl of each complex were applied to plasma-cleaned grids (Tergeo EM plasma Cleaner, PIE Scientific) which were blotted for 1.5-2.0 s using a Vitrobot Mark IV (Thermo Fisher Scientific) operated at 5°C and 100% humidity, and then plunged-frozen into liquid ethane cooled by liquid nitrogen. After vitrification, grids were clipped into autogrid cartridges (Thermo Fisher Scientific) to use with autoloader systems.

### Data acquisition

Grid screening and data collection were performed on a 200 kV Glacios cryo-transmission electron microscope equipped with a Falcon 4i direct electron detector (Thermo Fisher Scientific). Four grids for each complex were screened, and the best grid, as judged by ice thickness and particle distribution, was chosen for data collection. Grid squares were manually selected and automatically brought to eucentric height in EPU (Thermo Fisher Scientific), and holes were automatically detected and selected using specific relative ice thickness filters suitable for cyclin A-Cdk2 complexes. Electron micrograph movies were collected in EER format with a pixel size of 0.5675 Å/pixel, a total electron exposure of 60 e^-^/Å^2^, and a defocus range of -0.6 to -1.8 μm. In total, 6,422 movies were collected for Cdk2-cyclin A-GMNC, 5,078 for Cdk2-cyclin A-SAMHD1, and 4,707 for Cdk2-cyclin A-SCAPER.

### Image processing

For all structures, cryo-EM data were pre-processed in cryoSPARC live and cryoSPARC v4.4.1 (Punjani et al., 2017), and then transferred to RELION 5.0 beta^63^ for further processing, 3D reconstruction, and final refinement. All refinements were done using BLUSH regularization implemented in RELION 5.0 beta^63^.

In cryoSPARC live, raw EER movies were fractioned into 40 frames and motion corrected with 2x binning. The contrast transfer function (CTF) of each motion-corrected micrograph was fitted in cryoSPARC live. Total beam-induced specimen motion, in-frame motion, relative ice-thickness, and the quality of CTF fitting were inspected to remove poor-quality micrographs, which led to 6,015 micrographs being accepted for Cdk2-cyclin A-GMNC, 5,078 for Cdk2-cyclin A-SAMHD1 and 4,077 for Cdk2-cyclin A-SCAPER. For all three datasets, particles were picked using blob picker (circular/elliptical blob, 70-90 Å diameter) and then extracted using a box size of 160x160 pixels. On-the-fly 2D classification and 3D refinement using default parameters was used to assess data quality and peptide binding. Using a volume generated from the predicted structure of a Cdk2-cyclin A complex^64^ using UCSF Chimera^65^ as an initial reference, 447,768 particles yielded a 3.5 Å reconstruction for Cdk2-cyclin A-GMNC, 589,358 particles yielded a 3.3 Å reconstruction for Cdk2-cyclin A-SAMHD1, and 339,520 particles yielded a 3.4 Å reconstruction for Cdk2-cyclin A-SCAPER, according to Fourier Shell Correlation (FSC) at the 0.143 cut-off^66^.

To maximize the selection of high-quality particles, two particle selection strategies were employed in cryoSPARC v4.4.1^67^ for all three datasets after completion of on-the-fly processing. All blob-picked particles from the cryoSPARC live session were re-classified into 100 2D classes using a batch size of 300 particles, which resulted in 1,512,409 particles from good 2D classes retained for Cdk2-cyclin A-GMNC, 1,040,145 particles for Cdk2-cyclin A-SAMHD1, and 1,037,921 particles for Cdk2-cyclin A-SCAPER (particle set (i) identified in **Fig S3A**). Particles were also extracted using template picking using 2D averages generated by the reclassification of blob-picked particles from cryoSPARC live. Template-based picked particles were classified into 100 2D classes using a batch size of 300 particles which resulted in 1,280,980 particles from good 2D classes retained for Cdk2-cyclin A-GMNC, 916,334 particles for Cdk2-cyclin A-SAMHD1, and 869,030 particles for Cdk2-cyclin A-SCAPER (particle set (ii) identified in **Fig S3A**). Particles picked by blob-picking (particle set (i)) and template-picking (particle set (ii)) were combined with particles retained from good 2D classes from the cryoSPARC live session (particle set (iii) identified in **Fig S3A**). After removing duplicates, 2,553,964 particles were retained for Cdk2-cyclin A-GMNC, 1,712,010 for Cdk2-cyclin A-SAMHD1, and 1,741,162 for Cdk2-cyclin A-SCAPER. Retained particles were further classified into 50 2D classes using a batch size of 200 particles and duplicates were removed. After selecting good 2D classes, 1,938,556 particles were retained for Cdk2-cyclin A-GMNC, 1,473,643 for Cdk2-cyclin A-SAMHD1, and 1,446,486 for Cdk2-cyclin A-SCAPER, which were then converted to *.star files for use in RELION 5.0 beta using scripts contained in the PYEM package^68^.

Motion-corrected micrographs from CryoSPARC live were imported to RELION 5.0 beta and selected particles were re-extracted using a 160x160 pixel box using the coordinate information from the imported particles from cryoSPARC v4.4.1. Extracted particles were subjected to masked 3D refinement. Particles were then classified into four 3D classes by 3D classification without alignment (regularization parameter τ = 20). 183,712 particles for Cdk2-cyclin A-GMNC, 81,294 particles for Cdk2-cyclin A-SAMHD1, and 91,934 particles for Cdk2-cyclin A-SCAPER were selected from the best 3D class and refined to 3.6 Å resolution, 3.4 Å resolution, and 3.3 Å resolution respectively. To further improve the map quality, selected particles were subjected to CTF refinement (beam tilt and trefoil for the SAMHD1 and SCAPER datasets, and beam tilt, trefoil, and fourth-order aberrations for the GMNC dataset). For Cdk2-cyclin A-GMNC, CTF-refined particles were refined and post-processed yielding a 3.2 Å reconstruction, according to FSC at the 0.143 cut-off (**Fig S3B**). For Cdk2-cyclin A-SAMHD1 and Cdk2-cyclin A-SCAPER, CTF-refined particles were then re-refined and duplicates were removed, retaining 80,673 particles and 91,179 particles, respectively. The retained particles were then again refined and post-processed yielding a 3.1 Å reconstruction for Cdk2-cyclin A-SAMHD1 (**Fig S5A**) and a 2.9 Å reconstruction for CDK2-cyclin A-SCAPER (**Fig S6C**), according to FSC at the 0.143 cut-off.

### Model building and refinement

To build the peptide-bound Cdk2-cyclin A structures an AlphaFold-predicted model^64^ was fitted into each post-processed cryo-EM map (3.2 Å resolution for Cdk2-cyclin A-GMNC, 3.1 Å resolution for Cdk2-cyclin A-SAMHD1 and 2.9 Å resolution for Cdk2-cyclin A-SCAPER) using UCSF Chimera^65^. Fitted models were transferred to COOT for model building^69^ and refined by real-space refinement in PHENIX^70^. Refined models were validated using MOLPROBITY^71^ implemented in PHENIX. Refinement statistics for each structure are provided in **Table S8**. The three-dimensional FSC of each structure was computed using the Remote 3D FSC Processing server^72^.

### Visualization of molecular models and creation of figures

Molecular models and cryo-EM maps were visualized in UCSF Chimera^65^, UCSF Chimera X^73^, and PyMOL (the PyMOL molecular graphics system, Schrödinger, LLC) for analysis, interpretation and figure preparation.

---

**Figure S1.**
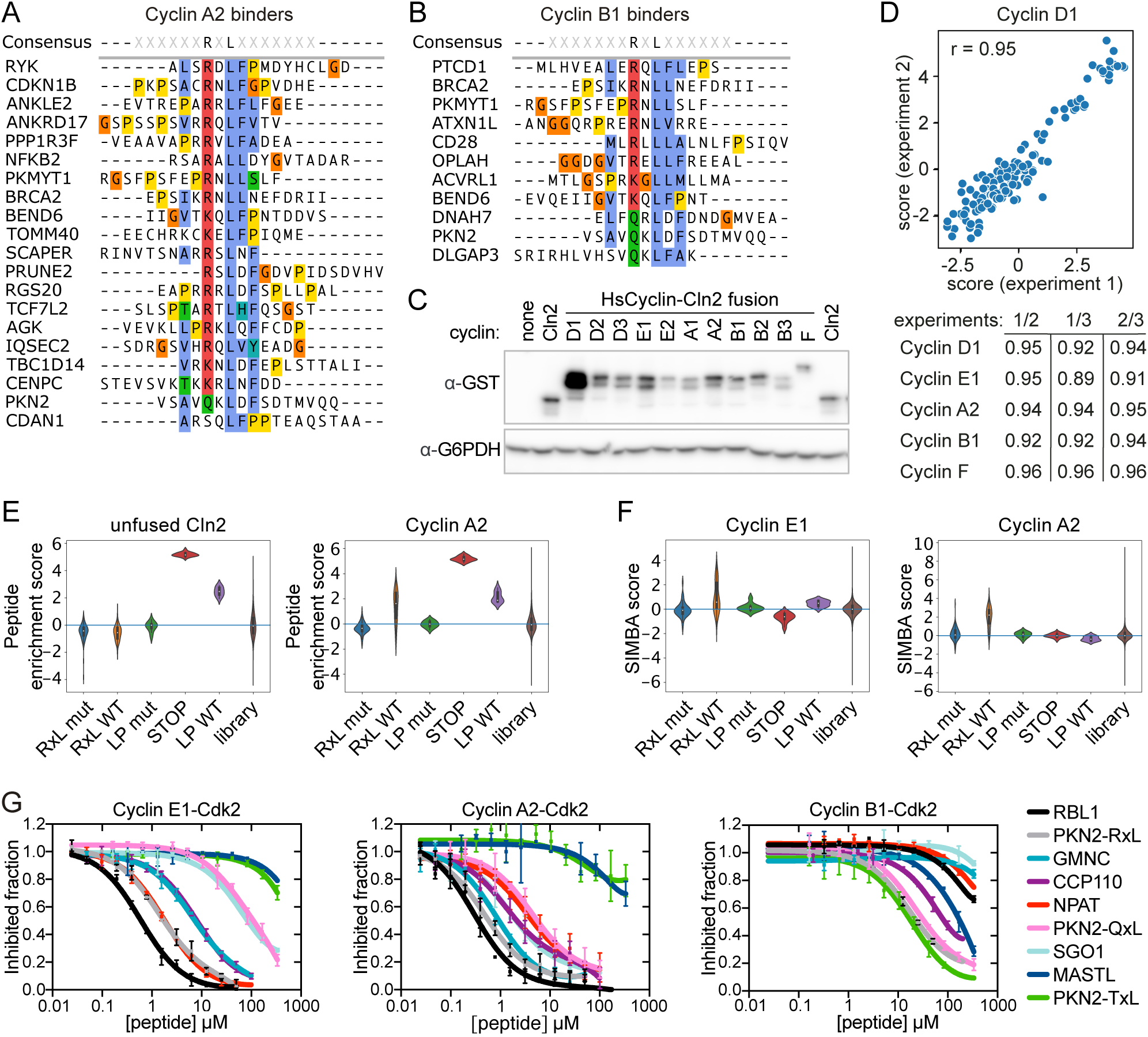
Characterizing cyclin binders with proteomic peptide phage display, SIMBA, and fluorescence polarization. (**A-B**) Alignment of RxL and RxL-like peptides identified as cyclin A2 (A) and B1 (B) binders in proteomic peptide phage display experiments. (**C**) The expression of 11 HsCyclin-Cln2 fusions was confirmed by Western blotting and probing for the GST tag. G6PDH serves as a loading control. (**D**) Correlation of peptide enrichment scores between three replicate SIMBA experiments screening the tiled peptide library. The correlation coefficients (r) were calculated from the control peptides. (**E**) Violin plots showing the enrichment scores of the indicated groups of peptides in the tiled library screen. (**F**) Violin plots of SIMBA scores, for different groups of peptides, in which the enrichment score of each peptide in HsCyclin-Cln2 strains is normalized by the score in the unfused Cln2 strain. (**G**) Dose-response curves showing the ability of various unlabeled competitor peptides to displace a FITC-labeled PKN2-RxL peptide from cyclin A2-, E1- and B1-Cdk2 complexes, measured by fluorescence polarization.

**Figure S2.**
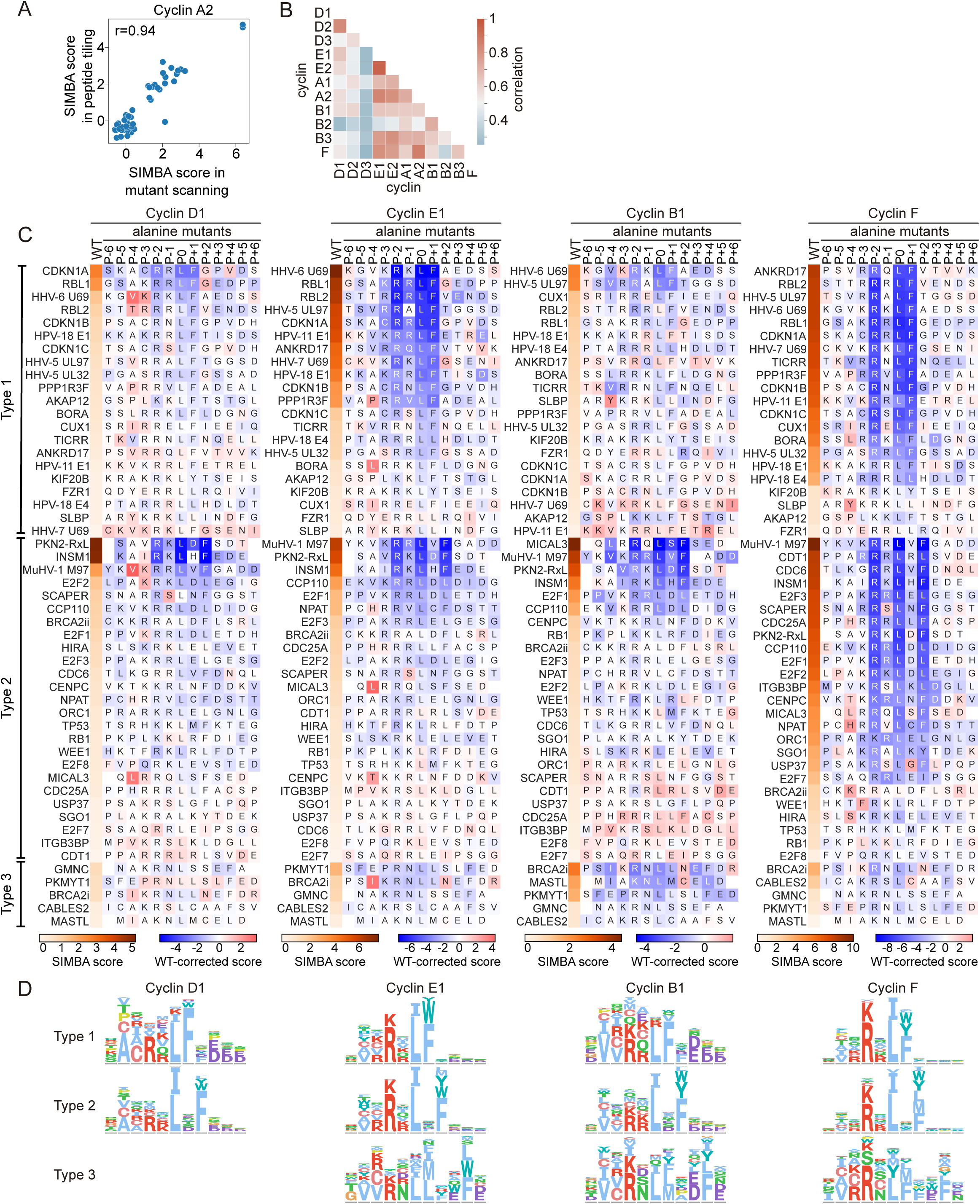
Uncovering the RxL binding determinants by mutational scanning in SIMBA. (**A**) Correlation of the control peptide SIMBA scores for cyclin A2 in cell cycle protein peptide tiling and mutant scanning SIMBA experiments. (**B**) Pairwise correlations between 11 cyclins showing the similarities in the SIMBA scores for the 7,858 distinct peptide variants in the mutational scanning library. (**C**) Heatmaps of RxL peptide alanine-scanning results for cyclins D1, E1, B1 and F, where the leftmost column shows the wild-type peptide SIMBA score, the letters show the peptide sequence and the color of the box shows the wild-type corrected score of the peptide with Ala at that position. (**D**) Amino acid preferences of different cyclins for type 1, 2 and 3 RxL motifs. The sequence logos show averaged preferences from DMS data of all starting peptides that bound the indicated cyclin.

**Figure S3.**
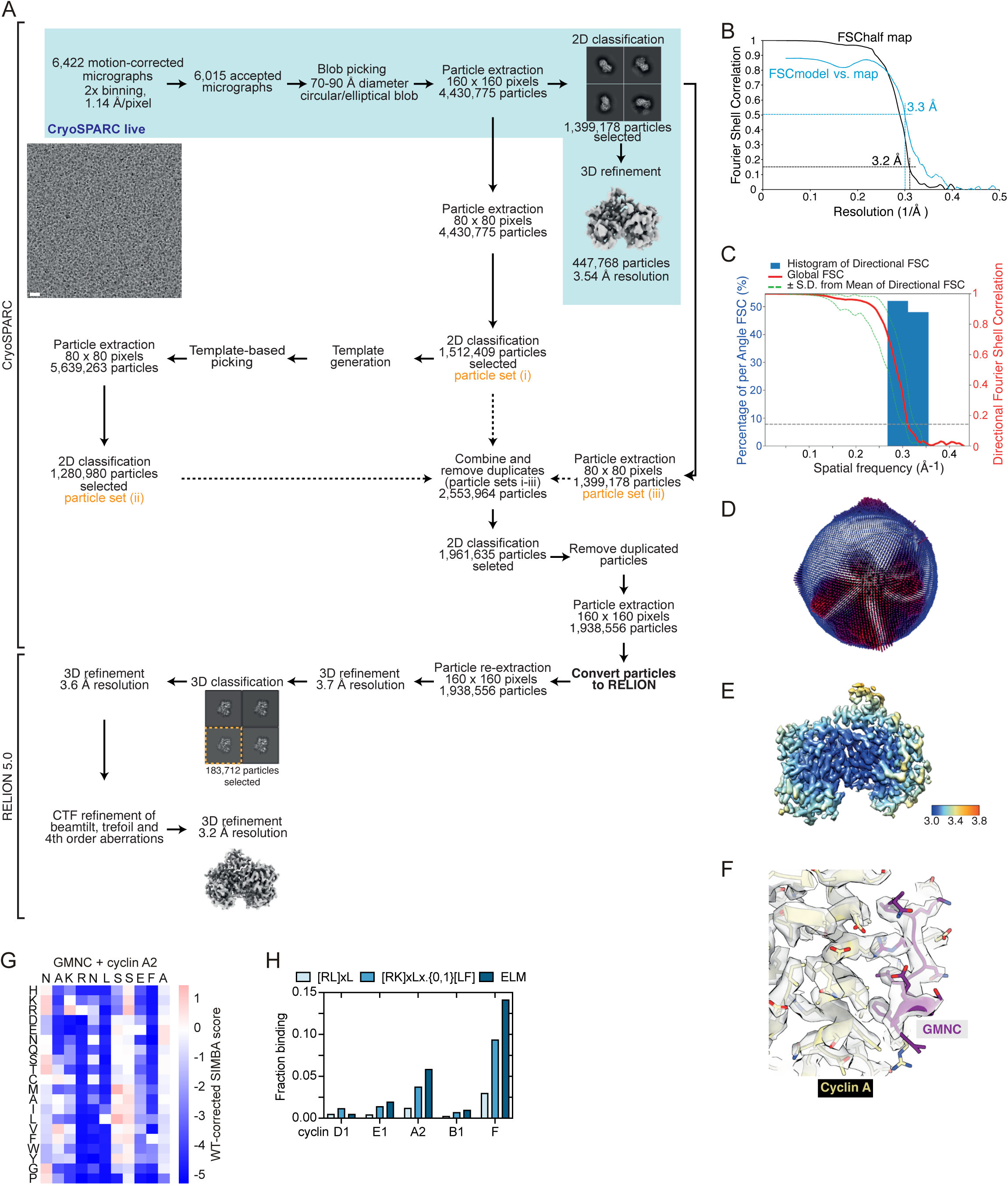
Structural characterization of GMNC type 3 RxL peptide interaction. **(A)** Data processing flowchart for Cdk2-cyclin A-GMNC RxL structure determination. 3D refined maps are shown as surface views, maps from 3D classification are shown as central slices. A representative micrograph is shown (scale bar: 150 Å). The micrograph was low-pass filtered to facilitate visualization of the particles. (**B-F**) Validation of the peptide-bound Cdk2-cyclin A-GMNC cryo-EM map. Validation metrics shown: **(B)** Fourier shell correlation (FSC) curves (half-map FSC in black, model vs. map FSC in blue), **(C)** analysis of the cryo-EM map using the 3D FSC server^72^, **(D)** orientation distribution plot, **(E)** local resolution estimates computed using RELION, and **(F)** section of the cryo-EM map with fitted models showing density for the peptides. (**G**) DMS results of GMNC RxL with cyclin A2. The heatmap shows wild-type corrected scores. (**H**) Plot showing the fraction of peptides matching different RxL consensus definitions that were identified as binders to the indicated cyclins in the tiled library screen.

**Figure S4.**
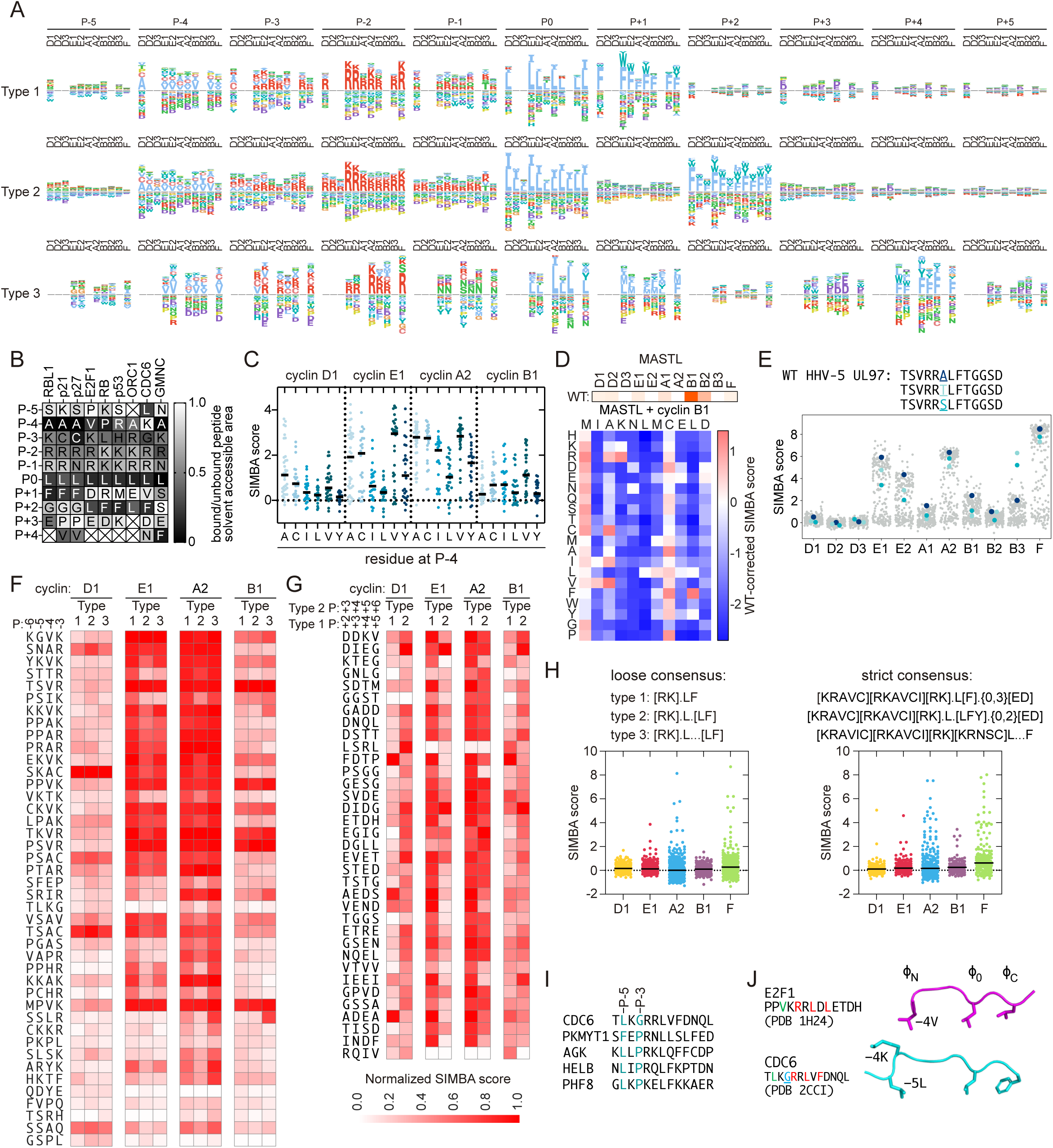
Mutational scanning identifies cyclin-specific preferences in RxL motifs. (**A**) Sequence logos showing the favored and disfavored residues at each position of RxL motifs for all tested cyclins. DMS data from peptides where the wild-type is a binder for the respective cyclin were averaged separately for each RxL type. Data is absent where a cyclin bound none of the wild-type peptides. (**B**) Relative solvent accessible area in bound RxL peptides compared to unbound peptides shows the buried residues in eight RxL-cyclin structures (1H24-28, 2CCI, 6P8H, 6P3W). (**C**) SIMBA scores of mutationally scanned RxL peptides with the indicated P-4 residues are shown for cyclins D1, E1, A2 and B1. Lines, median. (**D**) SIMBA scores of the wild-type MASTL RxL for each cyclin (top) and the DMS results for cyclin B1 plotted as wild-type corrected scores (bottom). (**E**) SIMBA scores of UL97 RxL DMS peptides for all tested cyclins, highlighting P-1 substitutions that increase B3 binding. (**F**) The impact of N-terminal flanking sequences on RxL binding was tested by joining 42 different N-terminal flanking sequences to the core sequences from single type 1, 2, or 3 RxL motifs, derived from RBL1 (RRLFGEDPP), CCP110 (RRLDLDIDG) and BRCA2 (RNLLNEFDR), respectively. Heatmaps show column-normalized SIMBA scores. (**G**) 35 C-terminal flanking sequences were joined to core sequences from type 1 (RBL1, GSAKRRLF) and type 2 (CCP110, EKVKRRLDL) motifs. Heatmaps show column-normalized SIMBA scores. (**H**) The SIMBA scores of tested RxL peptides that match a loose (left) or strict (right) RxL consensus definition. Lines, median. (**I**) Sequences of five outlier peptides from the comparison of PSSM-predicted scores with SIMBA scores (Fig 4J). (**J**) Isolated structures of cyclin-bound RxL peptides showing that the E2F1 P-4 Val and CDC6 P-5 Leu residues bind the same pocket on cyclin A2 (PDB: 1H24, 2CCI).

**Figure S5.**
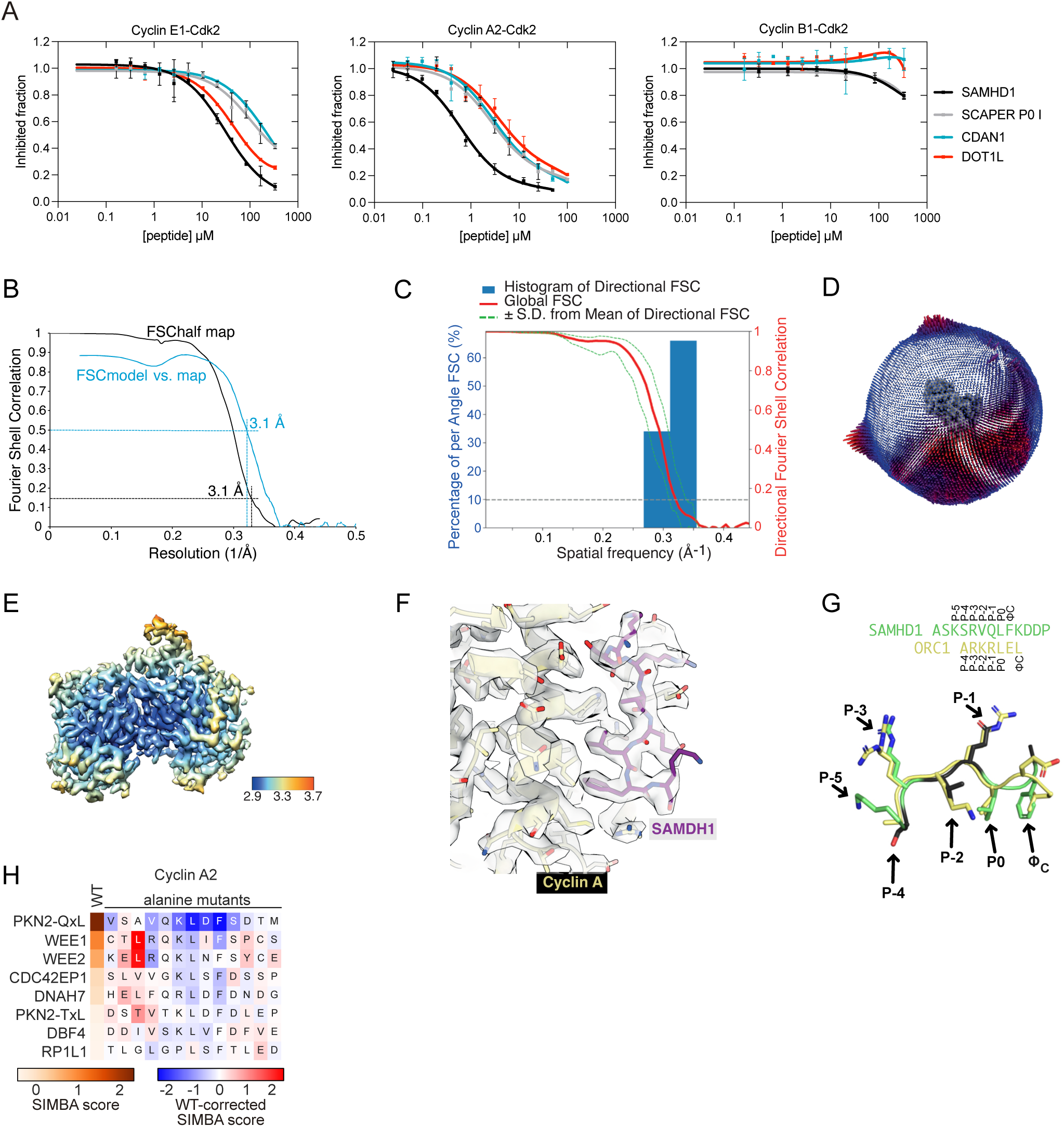
Characterization of atypical cyclin docking motifs lacking a P-2 basic residue. **(A)** Dose-response curves of FP assays show the concentrations of unlabeled competitor peptides needed to displace a FITC-labeled PKN2-RxL peptide from cyclin A2-, E1- and B1-Cdk2 complexes. (**B-F**) Validation of the peptide-bound Cdk2-cyclin A-SAMHD1 cryo-EM map. Validation metrics shown: **(B)** Fourier shell correlation (FSC) curves (half-map FSC in black, model vs. map FSC in blue), **(C)** analysis of the cryo-EM map using the 3D FSC server^72^, (D) orientation distribution plot, **(E)** local resolution estimates computed using RELION, and **(F)** section of the cryo-EM map with fitted models showing density for the peptides. (**G**) Overlay of ORC1 RxL (PDB 6P3W) and SAMHD1 RxL-like peptide in their cyclin-bound conformation shows similar binding conformation of the peptides despite the lack of a basic P-2 residue in SAMHD1. (**H**) SIMBA results of cyclin A2 binding to Ala-scanned LxF-type motifs. The plot shows wild-type-corrected scores for peptides with Ala at each position. The first column shows the SIMBA score of the wild-type peptide.

**Figure S6.**
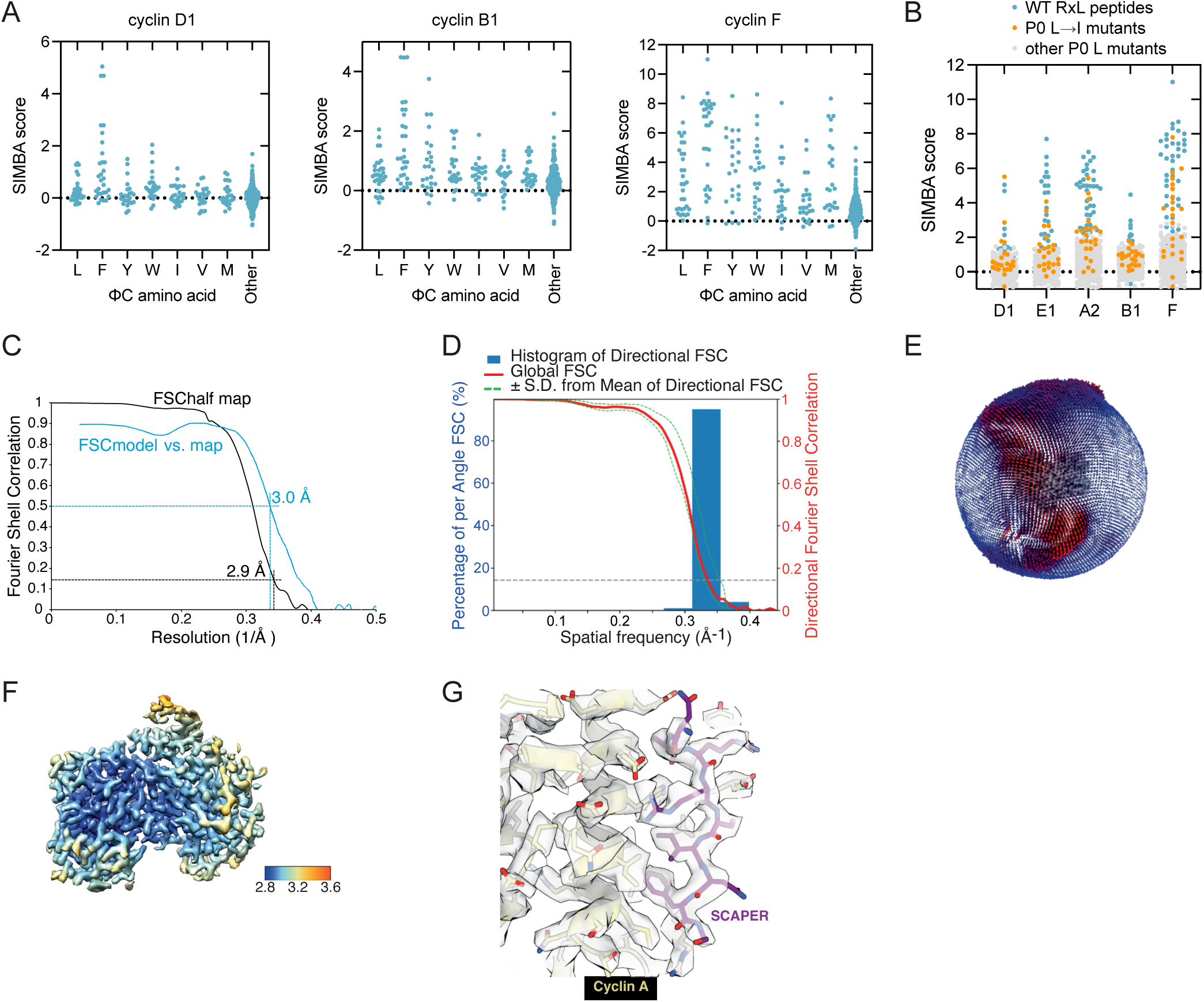
Tolerance for non-canonical residues at the Φ_0_ and Φ_C_ positions of cyclin docking motifs. (**A**) Results from DMS experiments showing SIMBA scores of RxL peptides with different residues at the Φ_C_ position. (**B**) Results from DMS experiments showing SIMBA scores of wild-type (WT) peptides and mutant peptides harboring Ile or other mutations at Φ_0_. (**C-G**) Validation of the peptide-bound Cdk2-cyclin A-SAMHD1 cryo-EM map. Validation metrics shown: **(C)** Fourier shell correlation (FSC) curves (half-map FSC in black, model vs. map FSC in blue), **(D)** analysis of the cryo-EM map using the 3D FSC server^72^, **(E)** orientation distribution plot, **(F)** local resolution estimates computed using RELION, and **(G)** section of the cryo-EM map with fitted models showing density for the peptides.

**Figure S7.**
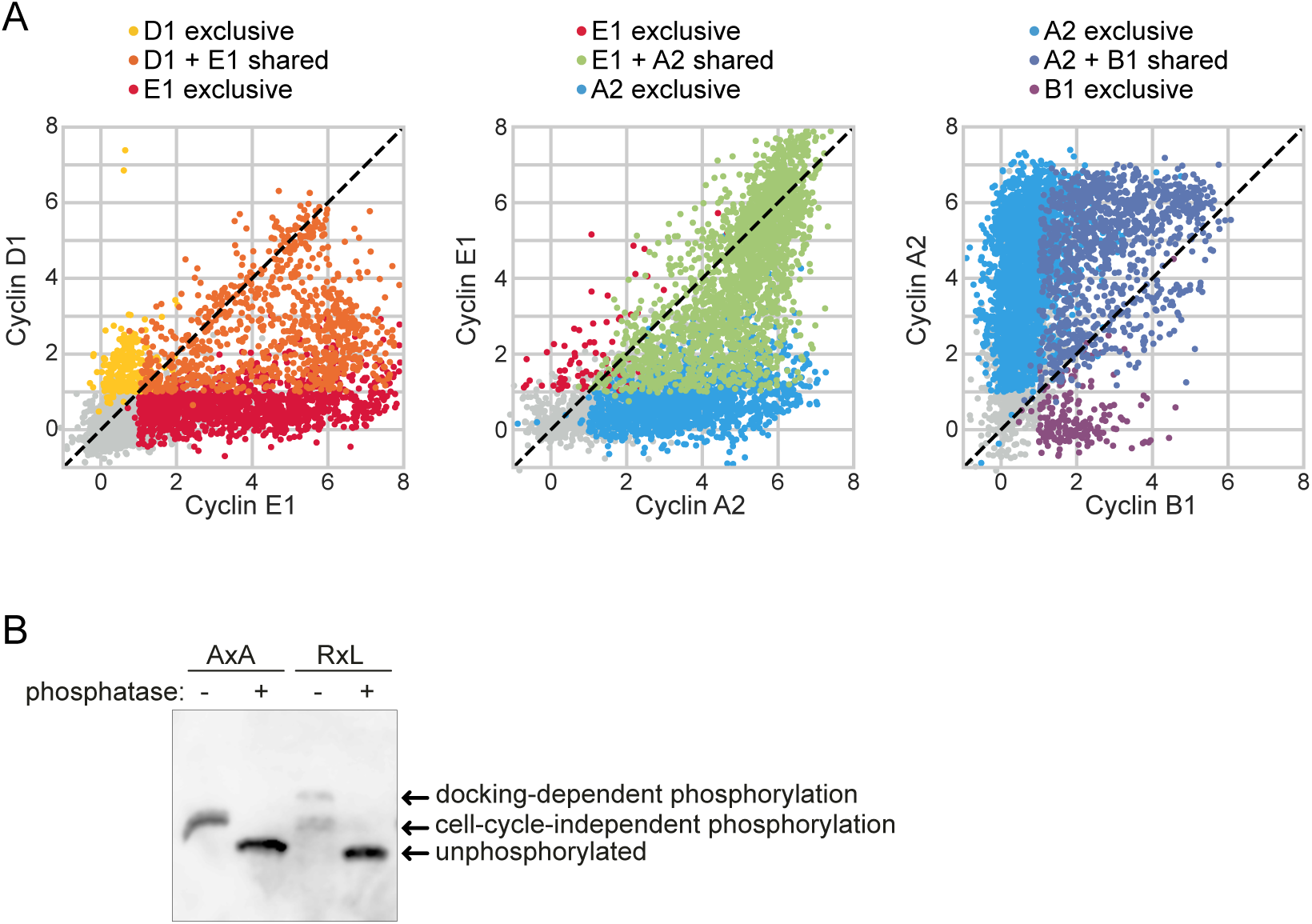
Repertoire of docking motifs recognized by cyclins during the cell cycle. (**A**) Pairwise comparisons of cyclins, ordered by their sequential expression, for SIMBA scores of mutationally scanned peptides illustrates the dynamics of cyclin docking specificity over the cell cycle. (**B**) Lysates from asynchronous RPE1 cultures expressing the PKMYT1-based reporter with either a control motif (AxA) or a PKN2-RxL docking motif (RxL) were treated with or without lambda phosphatase and analyzed for phosphorylation using Phos-tag SDS-PAGE Western blotting. A representative example of three experiments is shown. PKMYT1 reporters were detected using α-FLAG antibody.

## Notes

https://doi.org/10.17632/bs54ny7ns2.1

### Competing Interest Statement

The authors have declared no competing interest.

## References

1. Morgan, D.O. (2007). The cell cycle: principles of control (New Science Press).

2. Daub, H., Olsen, J.V., Bairlein, M., Gnad, F., Oppermann, F.S., Körner, R., Greff, Z., Kéri, G., Stemmann, O., and Mann, M. (2008). Kinase-selective enrichment enables quantitative phosphoproteomics of the kinome across the cell cycle. Mol. Cell 31, 438–448. 10.1016/J.MOLCEL.2008.07.007.

3. Dephoure, N., Zhou, C., Villén, J., Beausoleil, S.A., Bakalarski, C.E., Elledge, S.J., and Gygi, S.P. (2008). A quantitative atlas of mitotic phosphorylation. Proc. Natl. Acad. Sci. U. S. A. 105, 10762–10767. 10.1073/PNAS.0805139105.

4. Errico, A., Deshmukh, K., Tanaka, Y., Pozniakovsky, A., and Hunt, T. (2010). Identification of substrates for cyclin dependent kinases. Adv. Enzyme Regul. 50, 375–399. 10.1016/J.ADVENZREG.2009.12.001.

5. Pagliuca, F.W., Collins, M.O., Lichawska, A., Zegerman, P., Choudhary, J.S., and Pines, J. (2011). Quantitative Proteomics Reveals the Basis for the Biochemical Specificity of the Cell-Cycle Machinery. Mol. Cell 43, 406–417. 10.1016/j.molcel.2011.05.031.

6. Matthews, H.K., Bertoli, C., and Bruin, R.A.M. de (2021). Cell cycle control in cancer. Nat. Rev. Mol. Cell Biol. 2021 231 23, 74–88. 10.1038/s41580-021-00404-3.

7. Kõivomägi, M., Valk, E., Venta, R., Iofik, A., Lepiku, M., Morgan, D.O., and Loog, M. (2011). Dynamics of Cdk1 Substrate Specificity during the Cell Cycle. Mol. Cell 42, 610–623. 10.1016/j.molcel.2011.05.016.

8. Topacio, B.R., Zatulovskiy, E., Cristea, S., Xie, S., Tambo, C.S., Rubin, S.M., Sage, J., Koivomägi, M., and Skotheim, J.M. (2019). Cyclin D-Cdk4,6 Drives Cell-Cycle Progression via the Retinoblastoma Protein’s C-Terminal Helix. Mol. Cell 0. 10.1016/j.molcel.2019.03.020.

9. Holder, J., Poser, E., and Barr, F.A. (2019). Getting out of mitosis: spatial and temporal control of mitotic exit and cytokinesis by PP1 and PP2A. FEBS Lett. 593, 2908–2924. 10.1002/1873-3468.13595.

10. al-Rawi, A., Kaye, E., Korolchuk, S., Endicott, J.A., and Ly, T. (2023). Cyclin A and Cks1 promote kinase consensus switching to non-proline-directed CDK1 phosphorylation. Cell Rep. 42. 10.1016/J.CELREP.2023.112139.

11. Suzuki, K., Sako, K., Akiyama, K., Isoda, M., Senoo, C., Nakajo, N., and Sagata, N. (2015). Identification of non-Ser/Thr-Pro consensus motifs for Cdk1 and their roles in mitotic regulation of C2H2 zinc finger proteins and Ect2. Sci. Rep. 5, 7929. 10.1038/srep07929.

12. McGrath, D.A., Balog, E.R.M., Koivomägi, M., Lucena, R., Mai, M.V., Hirschi, A., Kellogg, D.R., Loog, M., and Rubin, S.M. (2013). Cks confers specificity to phosphorylation-dependent CDK signaling pathways. Nat. Struct. Mol. Biol. 20, 1407–1414. 10.1038/nsmb.2707.

13. Kõivomägi, M., Örd, M., Iofik, A., Valk, E., Venta, R., Faustova, I., Kivi, R., Balog, E.R.M., Rubin, S.M., and Loog, M. (2013). Multisite phosphorylation networks as signal processors for Cdk1. Nat. Struct. Mol. Biol. 20. 10.1038/nsmb.2706.

14. Veld, P.J.H. in ‘t, Wohlgemuth, S., Koerner, C., Müller, F., Janning, P., and Musacchio, A. (2022). Reconstitution and use of highly active human CDK1:Cyclin-B:CKS1 complexes. Protein Sci. Publ. Protein Soc. 31, 528–537. 10.1002/PRO.4233.

15. Tatum, N.J., and Endicott, J.A. (2020). Chatterboxes: the structural and functional diversity of cyclins. Semin. Cell Dev. Biol. 10.1016/j.semcdb.2020.04.021.

16. Schulman, B.A., Lindstrom, D.L., and Harlow, E.D. (1998). Substrate recruitment to cyclin-dependent kinase 2 by a multipurpose docking site on cyclin A. Biochemistry 95, 10453–10458.

17. Wohlschlegel, J.A., Dwyer, B.T., Takeda, D.Y., and Dutta, A. (2001). Mutational analysis of the Cy motif from p21 reveals sequence degeneracy and specificity for different cyclin-dependent kinases. Mol. Cell. Biol. 21, 4868–4874. 10.1128/MCB.21.15.4868-4874.2001.

18. Lacy, E.R., Filippov, I., Lewis, W.S., Otieno, S., Xiao, L., Weiss, S., Hengst, L., and Kriwacki, R.W. (2004). p27 binds cyclin–CDK complexes through a sequential mechanism involving binding-induced protein folding. Nat. Struct. Mol. Biol. 2004 114 11, 358–364. 10.1038/nsmb746.

19. Takeda, D.Y., Wohlschlegel, J.A., and Dutta, A. (2001). A bipartite substrate recognition motif for cyclin-dependent kinases. J. Biol. Chem. 276, 1993–1997. 10.1074/jbc.M005719200.

20. Bailly, E., Cabantous, S., Sondaz, D., Bernadac, A., and Simon, M.-N. (2003). Differential cellular localization among mitotic cyclins from Saccharomyces cerevisiae: a new role for the axial budding protein Bud3 in targeting Clb2 to the mother-bud neck. J. Cell Sci. 116, 4119– 4130. 10.1242/jcs.00706.

21. Basu, S., Roberts, E.L., Jones, A.W., Swaffer, M.P., Snijders, A.P., and Nurse, P. (2020). The Hydrophobic Patch Directs Cyclin B to Centrosomes to Promote Global CDK Phosphorylation at Mitosis. Curr. Biol. 0. 10.1016/j.cub.2019.12.053.

22. Bentley, A.M., Normand, G., Hoyt, J., and King, R.W. (2007). Distinct sequence elements of cyclin B1 promote localization to chromatin, centrosomes, and kinetochores during mitosis. Mol. Biol. Cell 18, 4847–4858. 10.1091/mbc.E06-06-0539.

23. Dangiolella, V., Donato, V., Vijayakumar, S., Saraf, A., Florens, L., Washburn, M.P., Dynlacht, B., and Pagano, M. (2010). SCFCyclin F controls centrosome homeostasis and mitotic fidelity via CP110 degradation. Nature 466, 138. 10.1038/NATURE09140.

24. Allan, L.A., Reis, M.C., Ciossani, G., Veld, P.J.H. in ‘t, Wohlgemuth, S., Kops, G.J., Musacchio, A., and Saurin, A.T. (2020). Cyclin B1 scaffolds MAD1 at the kinetochore corona to activate the mitotic checkpoint. EMBO J. 39. 10.15252/EMBJ.2019103180.

25. Yu, J., Raia, P., Ghent, C.M., Raisch, T., Sadian, Y., Cavadini, S., Sabale, P.M., Barford, D., Raunser, S., Morgan, D.O., et al. (2021). Structural basis of human separase regulation by securin and CDK1–cyclin B1. Nat. 2021, 1–5. 10.1038/s41586-021-03764-0.

26. Jackman, M., Marcozzi, C., Barbiero, M., Pardo, M., Yu, L., Tyson, A.L., Choudhary, J.S., and Pines, J. (2020). Cyclin B1-Cdk1 facilitates MAD1 release from the nuclear pore to ensure a robust spindle checkpoint. J. Cell Biol. 219. 10.1083/jcb.201907082.

27. Bhaduri, S., and Pryciak, P.M. (2011). Cyclin-specific docking motifs promote phosphorylation of yeast signaling proteins by G1/S Cdk complexes. Curr. Biol. 21, 1615–1623. 10.1016/j.cub.2011.08.033.

28. Faustova, I., Bulatovic, L., Matiyevskaya, F., Valk, E., Örd, M., and Loog, M. (2020). A new linear cyclin docking motif that mediates exclusively S-phase CDK-specific signaling. EMBO J. 10.15252/embj.2020105839.

29. Örd, M., Venta, R., Möll, K., Valk, E., and Loog, M. (2019). Cyclin-Specific Docking Mechanisms Reveal the Complexity of M-CDK Function in the Cell Cycle. Mol. Cell 75, 76–89.e3. 10.1016/j.molcel.2019.04.026.

30. Örd, M., Puss, K.K., Kivi, R., Möll, K., Ojala, T., Borovko, I., Faustova, I., Venta, R., Valk, E., Koivomägi, M., et al. (2020). Proline-Rich Motifs Control G2-CDK Target Phosphorylation and Priming an Anchoring Protein for Polo Kinase Localization. Cell Rep. 31. 10.1016/j.celrep.2020.107757.

31. Cheng, K.Y., Noble, M.E.M., Skamnaki, V., Brown, N.R., Lowe, E.D., Kontogiannis, L., Shen, K., Cole, P.A., Siligardi, G., and Johnson, L.N. (2006). The role of the phospho-CDK2/cyclin A recruitment site in substrate recognition. J. Biol. Chem. 281, 23167–23179. 10.1074/jbc.M600480200.

32. Guiley, K.Z., Stevenson, J.W., Lou, K., Barkovich, K.J., Kumarasamy, V., Wijeratne, T.U., Bunch, K.L., Tripathi, S., Knudsen, E.S., Witkiewicz, A.K., et al. (2019). P27 allosterically activates cyclin-dependent kinase 4 and antagonizes palbociclib inhibition. Science 366. 10.1126/SCIENCE.AAW2106/SUPPL_FILE/AAW2106_GUILEY_SM.PDF.

33. Lowe, E.D., Tews, I., Cheng, K.Y., Brown, N.R., Gul, S., Noble, M.E.M., Gamblin, S.J., and Johnson, L.N. (2002). Specificity Determinants of Recruitment Peptides Bound to Phospho-CDK2/Cyclin A ^†^, ^‡^. Biochemistry 41, 15625–15634. 10.1021/bi0268910.

34. Gelais, C.S., Kim, S.H., Ding, L., Yount, J.S., Ivanov, D., Spearman, P., and Wu, L. (2016). A Putative Cyclin-binding Motif in Human SAMHD1 Contributes to Protein Phosphorylation, Localization, and Stability. J. Biol. Chem. 291, 26332. 10.1074/JBC.M116.753947.

35. D’Angiolella, V., Donato, V., Forrester, F.M., Jeong, Y.-T., Pellacani, C., Kudo, Y., Saraf, A., Florens, L., Washburn, M.P., and Pagano, M. (2012). Cyclin F-Mediated Degradation of Ribonucleotide Reductase M2 Controls Genome Integrity and DNA Repair. Cell 149, 1023– 1034. 10.1016/j.cell.2012.03.043.

36. Kelso, S., Orlicky, S., Beenstock, J., Ceccarelli, D.F., Kurinov, I., Gish, G., and Sicheri, F. (2021). Bipartite binding of the N terminus of Skp2 to cyclin A. Structure 29, 975–988.e5. 10.1016/J.STR.2021.04.011.

37. Salamina, M., Montefiore, B.C., Liu, M., Wood, D.J., Heath, R., Ault, J.R., Wang, L.Z., Korolchuk, S., Baslé, A., Pastok, M.W., et al. (2021). Discriminative SKP2 Interactions with CDK-Cyclin Complexes Support a Cyclin A-Specific Role in p27KIP1 Degradation. J. Mol. Biol. 433. 10.1016/J.JMB.2020.166795.

38. Singh, S., Gleason, C.E., Fang, M., Laimon, Y.N., Khivansara, V., Xie, S., Durmaz, Y.T., Sarkar, A., Ngo, K., Savla, V., et al. (2024). Cyclin A/B RxL Macrocyclic Inhibitors to Treat Cancers with High E2F Activity. BioRxiv Prepr. Serv. Biol., 2024.08.01.605889. 10.1101/2024.08.01.605889.

39. Hope, I., Noble, M.E.M., Waring, M.J., Noble, E.M.M., Endicott, J.A., and Tatum, N.J. (2024). Crystallographic fragment screening of CDK2-cyclin A: FragLites map sites of protein-protein interaction. Preprint at bioRxiv, 10.1101/2024.06.03.596235.

40. Subbanna, M.S., Winters, M.J., Örd, M., Davey, N.E., and Pryciak, P.M. (2024). A quantitative intracellular peptide binding assay reveals recognition determinants and context dependence of short linear motifs. Preprint at bioRxiv, 10.1101/2024.10.30.621084.

41. Ferrell, J.E. (2011). Simple Rules for Complex Processes: New Lessons from the Budding Yeast Cell Cycle. Mol. Cell 43, 497. 10.1016/J.MOLCEL.2011.08.002.

42. Zhang, T., Liu, W.D., Saunee, N.A., Breslin, M.B., and Lan, M.S. (2009). Zinc Finger Transcription Factor INSM1 Interrupts Cyclin D1 and CDK4 Binding and Induces Cell Cycle Arrest. J. Biol. Chem. 284, 5574. 10.1074/JBC.M808843200.

43. Hara, M., Abe, Y., Tanaka, T., Yamamoto, T., Okumura, E., and Kishimoto, T. (2012). Greatwall kinase and cyclin B-Cdk1 are both critical constituents of M-phase-promoting factor. Nat. Commun. 2012 31 3, 1–9. 10.1038/ncomms2062.

44. Balestrini, A., Cosentino, C., Errico, A., Garner, E., and Costanzo, V. (2010). GEMC1 is a TopBP1-interacting protein required for chromosomal DNA replication. Nat. Cell Biol. 2010 125 12, 484–491. 10.1038/ncb2050.

45. Liu, F., Rothblum-Oviatt, C., Ryan, C.E., and Piwnica-Worms, H. (1999). Overproduction of Human Myt1 Kinase Induces a G2 Cell Cycle Delay by Interfering with the Intracellular Trafficking of Cdc2-Cyclin B1 Complexes. Mol. Cell. Biol. 19, 5113. 10.1128/MCB.19.7.5113.

46. Brown, N.R., Lowe, E.D., Petri, E., Skamnaki, V., Antrobus, R., Johnson, L., and Johnson, L.N. (2007). Cyclin B and Cyclin A Confer Different Substrate Recognition Properties on CDK2. Cell Cycle 6, 11. 10.4161/cc.6.11.4278.

47. Kumar, M., Michael, S., Alvarado-Valverde, J., Sign©száros, B.M.D., Sámano-Sánchez, H., Zeke, A., Dobson, L., Lazar, T., Örd, M., Nagpal, A., et al. (2022). The Eukaryotic Linear Motif resource: 2022 release. Nucleic Acids Res. 50, D497. 10.1093/NAR/GKAB975.

48. Doig, A.J., and Baldwin, R.L. (1995). N- and C-capping preferences for all 20 amino acids in α-helical peptides. Protein Sci. 4, 1325–1336. 10.1002/pro.5560040708.

49. Chi, Y., Carter, J.H., Swanger, J., Mazin, A.V., Moritz, R.L., and Clurman, B.E. (2020). A novel landscape of nuclear human CDK2 substrates revealed by in situ phosphorylation. Sci. Adv. 6, 9899–9916. 10.1126/SCIADV.AAZ9899/SUPPL_FILE/AAZ9899_TABLE_S5.XLSX.

50. Trotter, E.W., and Hagan, I.M. (2020). Release from cell cycle arrest with Cdk4/6 inhibitors generates highly synchronized cell cycle progression in human cell culture: Cdk4/6 Induction Synchronisation. Open Biol. 10. 10.1098/RSOB.200200/.

51. Castro, A., and Lorca, T. (2018). Greatwall kinase at a glance. J. Cell Sci. 131, jcs222364. 10.1242/jcs.222364.

52. Bandyopadhyay, S., Bhaduri, S., Örd, M., Davey, N.E., Loog, M., and Pryciak, P.M. (2020). Comprehensive Analysis of G1 Cyclin Docking Motif Sequences that Control CDK Regulatory Potency In Vivo. Curr. Biol. 30, 4454–4466.e5. 10.1016/j.cub.2020.08.099.

53. Gavet, O., and Pines, J. (2010). Progressive Activation of CyclinB1-Cdk1 Coordinates Entry to Mitosis. Dev. Cell 18, 533–543. 10.1016/j.devcel.2010.02.013.

54. Örd, M., Möll, K., Agerova, A., Kivi, R., Faustova, I., Venta, R., Valk, E., and Loog, M. (2019). Multisite phosphorylation code of CDK. Nat. Struct. Mol. Biol. 26, 649–658. 10.1038/s41594-019-0256-4.

55. Basu, S., Greenwood, J., Jones, A.W., and Nurse, P. (2022). Core control principles of the eukaryotic cell cycle. Nat. 2022 6077918 607, 381–386. 10.1038/s41586-022-04798-8.

56. Benz, C., Ali, M., Krystkowiak, I., Simonetti, L., Sayadi, A., Mihalic, F., Kliche, J., Andersson, E., Jemth, P., Davey, N.E., et al. (2022). Proteome-scale mapping of binding sites in the unstructured regions of the human proteome. Mol. Syst. Biol. 18, e10584. 10.15252/MSB.202110584.

57. Jumper, J., Evans, R., Pritzel, A., Green, T., Figurnov, M., Ronneberger, O., Tunyasuvunakool, K., Bates, R., Žídek, A., Potapenko, A., et al. (2021). Highly accurate protein structure prediction with AlphaFold. Nat. 2021 5967873 596, 583–589. 10.1038/s41586-021-03819-2.

58. Cross, F.R., and Jacobson, M.D. (2000). Conservation and Function of a Potential Substrate-Binding Domain in the Yeast Clb5 B-Type Cyclin Downloaded from. Mol. Cell. Biol. 20, 4782–4790.

59. Winters, M.J., and Pryciak, P.M. (2018). Analysis of the thresholds for transcriptional activation by the yeast MAP kinases Fus3 and Kss1. Mol. Biol. Cell 29, 669–682. 10.1091/mbc.E17-10-0578.

60. Bravo, A.M., Typas, A., and Veening, J.W. (2022). 2FAST2Q: a general-purpose sequence search and counting program for FASTQ files. PeerJ 10. 10.7717/PEERJ.14041.

61. Örd, M., and Loog, M. (2020). Detection of multisite phosphorylation of intrinsically disordered proteins using phos-tag SDS-PAGE. In Methods in Molecular Biology (Humana Press Inc.), pp. 779–792. 10.1007/978-1-0716-0524-0_40.

62. Albanese, S.K., Parton, D.L., Işik, M., Rodríguez-Laureano, L., Hanson, S.M., Behr, J.M., Gradia, S., Jeans, C., Levinson, N.M., Seeliger, M.A., et al. (2018). An open library of human kinase domain constructs for automated bacterial expression. Biochemistry 57, 4675–4689. 10.1021/acs.biochem.7b01081.

63. Kimanius, D., Jamali, K., Wilkinson, M.E., Lövestam, S., Velazhahan, V., Nakane, T., and Scheres, S.H.W. (2024). Data-driven regularization lowers the size barrier of cryo-EM structure determination. Nat. Methods 21, 1216–1221. 10.1038/s41592-024-02304-8.

64. Abramson, J., Adler, J., Dunger, J., Evans, R., Green, T., Pritzel, A., Ronneberger, O., Willmore, L., Ballard, A.J., Bambrick, J., et al. (2024). Accurate structure prediction of biomolecular interactions with AlphaFold 3. Nature 630, 493–500. 10.1038/s41586-024-07487-w.

65. Pettersen, E.F., Goddard, T.D., Huang, C.C., Couch, G.S., Greenblatt, D.M., Meng, E.C., and Ferrin, T.E. (2004). UCSF Chimera – A visualization system for exploratory research and analysis. J. Comput. Chem. 25, 1605–1612. 10.1002/jcc.20084.

66. Rosenthal, P.B., and Henderson, R. (2003). Optimal determination of particle orientation, absolute hand, and contrast loss in single-particle electron cryomicroscopy. J. Mol. Biol. 333, 721–745. 10.1016/j.jmb.2003.07.013.

67. Punjani, A., Rubinstein, J.L., Fleet, D.J., and Brubaker, M.A. (2017). cryoSPARC: algorithms for rapid unsupervised cryo-EM structure determination. Nat. Methods 14, 290–296. 10.1038/nmeth.4169.

68. Asarnow, D., Palovcak, E., and Cheng, Y. (2019). asarnow/pyem: UCSF pyem v0.5. Version v0.5. Zenodo. 10.5281/zenodo.3576630.

69. Emsley, P., Lohkamp, B., Scott, W.G., and Cowtan, K. (2010). Features and development of Coot. Acta Crystallogr. D Biol. Crystallogr. 66, 486–501. 10.1107/S0907444910007493.

70. Afonine, P.V., Poon, B.K., Read, R.J., Sobolev, O.V., Terwilliger, T.C., Urzhumtsev, A., and Adams, P.D. (2018). Real-space refinement in PHENIX for cryo-EM and crystallography. Acta Crystallogr. Sect. Struct. Biol. 74, 531–544. 10.1107/S2059798318006551.

71. Williams, C.J., Headd, J.J., Moriarty, N.W., Prisant, M.G., Videau, L.L., Deis, L.N., Verma, V., Keedy, D.A., Hintze, B.J., Chen, V.B., et al. (2017). MolProbity: More and better reference data for improved all-atom structure validation. Protein Sci. 27, 293–315. 10.1002/pro.3330.

72. Tan, Y.Z., Baldwin, P.R., Davis, J.H., Williamson, J.R., Potter, C.S., Carragher, B., and Lyumkis, D. (2017). Addressing preferred specimen orientation in single-particle cryo-EM through tilting. Nat. Methods 14, 793–796. 10.1038/nmeth.4347.

73. Goddard, T.D., Huang, C.C., Meng, E.C., Pettersen, E.F., Couch, G.S., Morris, J.H., and Ferrin, T.E. (2018). UCSF ChimeraX: Meeting modern challenges in visualization and analysis. Protein Sci. Publ. Protein Soc. 27, 14–25. 10.1002/pro.3235.

